# A novel system to map protein interactions reveals evolutionarily conserved immune evasion pathways on transmissible cancers

**DOI:** 10.1101/831404

**Authors:** Andrew S. Flies, Jocelyn M. Darby, Patrick R. Lennard, Peter R. Murphy, Chrissie E. B. Ong, Terry L. Pinfold, A. Bruce Lyons, Gregory M. Woods, Amanda L. Patchett

## Abstract

Immune checkpoint immunotherapy has revolutionized medicine, but translational success for new treatments remains low. Around 40% of humans and Tasmanian devils (*Sarcophilus harrisii*) develop cancer in their lifetime, compared to less than 10% for most species. Additionally, devils are affected by two of the three known transmissible cancers in mammals. Unfortunately, little is known about of immune checkpoints in devils and other non-model species, largely due to a lack of species-specific reagents. We developed a simple cut-and-paste reagent development method applicable to any vertebrate species and show that immune checkpoint interactions are conserved across 160 million years of evolution. The inhibitory checkpoint molecule CD200 is highly expressed on devil facial tumor cells. We are the first to demonstrate that co-expression of CD200R1 can block CD200 expression. The evolutionarily conserved pathways suggest that naturally occurring cancers in devils and other species can serve as models for understanding cancer and immunological tolerance.

**GRAPHICAL ABSTRACT:** 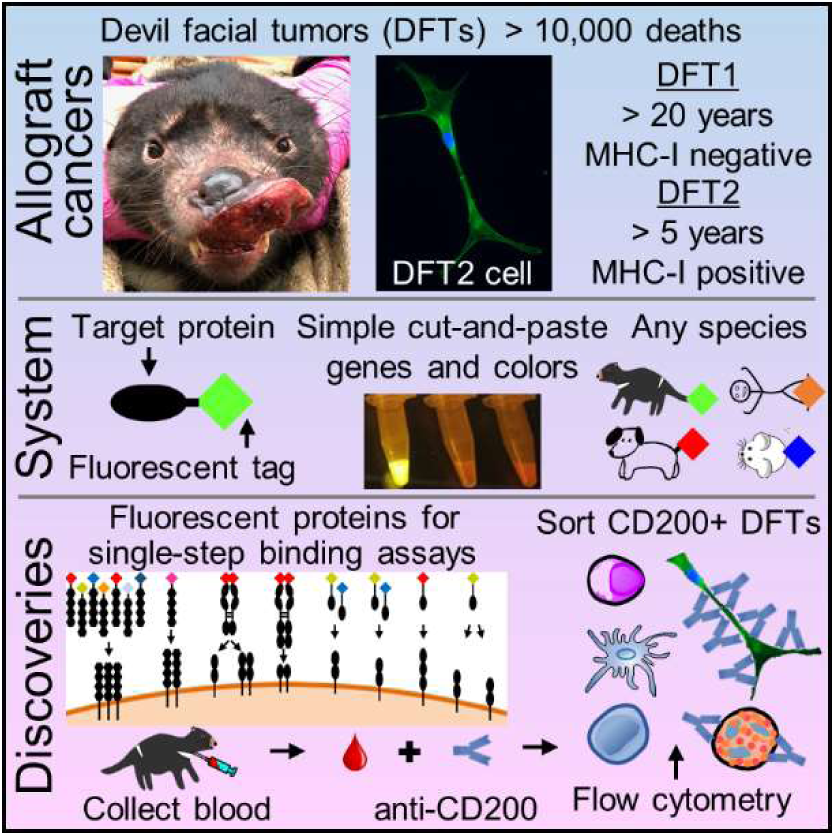

## INTRODUCTION

Metastatic cancer affects most mammals, but the cancer incidence can vary widely across phylogenetic groups and species (Figure 1, **Table S1**) ^1–9^. In humans, the lifetime risk of developing cancer is around 40% ^10^. This is in stark contrast to a general cancer incidence of 3% for mammals, 2% for birds, and 2% for reptiles reported by the San Diego Zoo (n=10,317) ^7, 11^. A more recent study at the Taipei Zoo reported cancer incidence of 8%, 4%, and 1% for mammals, birds, and reptiles, respectively (n=2,657) ^4^. Cancer incidence in domestic animals is generally less than 10% (n=202,277) ^9^. However, two studies performed 40 years apart reported that greater than 40% of Tasmanian devils develop spontaneous, often severe neoplasia in their lifetime ^11, 12^. Devils are also unique because they are affected by two of the three known naturally-occurring transmissible cancers in vertebrate species ^13, 14^. Transmissible cancers are a distinct form of cancer in which the tumor cells function as an infectious pathogen and an allograft. Dogs (*Canis lupus familiaris*) are the only other vertebrate species affected by a transmissible cancer ^15^, and interestingly some breeds of dogs also have high cancer incidence ^9, 16^.

**Figure 1.**
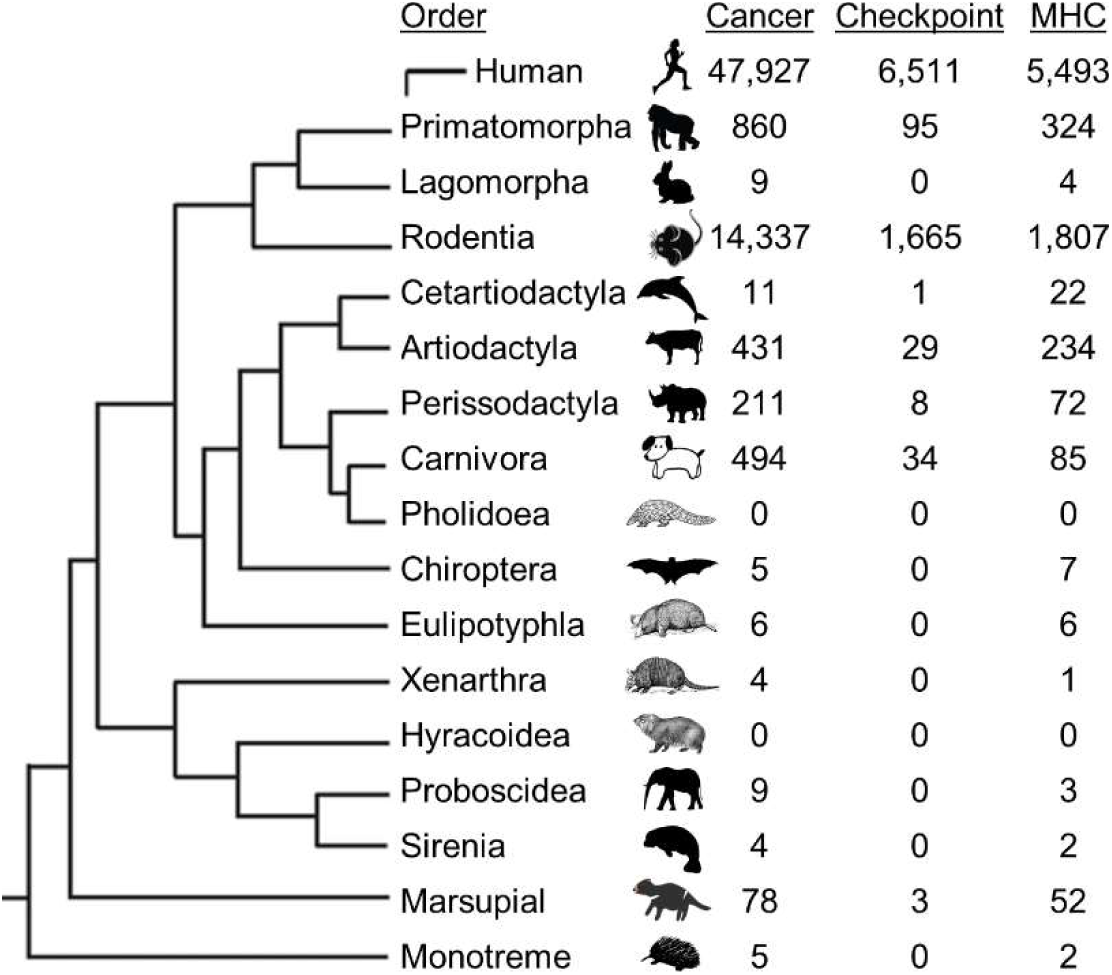
Phylogenetic tree of immune system related studies in mammal orders 2009-2019. Metastatic cancer has been reported in nearly all mammalian orders and major histocompatibility complexes (MHC) have been the most intensely studied molecules in most orders. In the past decade, studies of immune checkpoint molecules (PD1, PDL1, CTLA4) have become a primary focus in humans and rodents. However, immune checkpoint studies in other species are limited, particularly at the protein level, due to the lack of species-specific reagents. This creates a vast gap in our understanding of the evolution of the mammalian immune system. The numbers in the columns represent the number studies matching Web of Science search results between 2009-2019. See **Table S1** for search terms.

The devil facial tumor (DFT) disease was first detected in Northwest Tasmania and has been a primary driver of an 80% decline in the wild Tasmanian devil population ^13, 17^. The clonal devil facial tumor (DFT1) cells have been continually transmitted among devils and is estimated to have killed at least 10,000 individuals since at least 1996. In 2014 a second independent transmissible Tasmanian devil facial tumor (DFT2) was discovered in wild devils ^14^ and 23 cases have been reported to date ^18^. Genetic mismatches, particularly in the major histocompatibility complex (MHC) genes should lead to rejection of these transmissible tumors. Consequently, the role of devil MHC has been a focus of numerous studies (Figure 1, **Table S1**) to understand the lack of rejection of the transmissible tumors. These studies have revealed that the DFT1 cells downregulate MHC class I (MHC-I) expression ^19^, a phenomenon observed in many human cancers ^20^. In contrast to DFT1 cells, the DFT2 cells do express MHC-I ^21^. DFT1 and DFT2 cells also have 2,884 and 3,591 single nucleotide variants, respectively, that are not present in 46 normal devil genomes ^22^. The continual transmission of DFT1 and DFT2, despite MHC-I expression by DFT2 cells and genetic mismatches between host and tumor, suggests that additional pathways are likely involved in immune evasion.

Human cancer treatment has been revolutionized in the past decade by manipulating interactions among immune checkpoint molecules ^23, 24^. These have proven broadly effective in part because they function across many different MHC types and tumor mutational patterns. However, these pathways have received little attention in transmissible cancers and other naturally occurring cancers in non-model species (Figure 1, **Table S1**) ^25–27^. We have previously shown that the inhibitory immune checkpoint molecule programmed death ligand 1 (PDL1) is expressed in the DFT microenvironment and is upregulated by interferon-gamma (IFN*γ*) *in vitro* ^25^. This finding led us to question which other immune checkpoint molecules play a role in immune evasion by the transmissible cancers and the devil’s high spontaneous cancer incidence. Understanding this immune evasion in a natural environment has the potential to help protect this endangered species and identify protein interactions that are conserved across divergent species to improve translational success of animal models ^27^. Unfortunately, a persistent limitation for immunology in non-traditional study species is a lack of species-specific reagents. Wildlife biologists and veterinarians are at the front lines of emerging infectious disease outbreaks, but they often lack species-specific reagents to fulfil the World Health Organization’s call for “cross-cutting R&D preparedness” and perform mechanistic immunological investigations ^28^.

To solve the paucity of reagents available for Tasmanian devils and address ongoing limitations for non-traditional study species, we developed a Fluorescent Adaptable Simple Theranostic (FAST) protein system that builds upon the diverse uses of fluorescent proteins previously reported ^29–33^. This simple system can be used for rapid development of diagnostic and therapeutic (i.e. theranostic) immunological toolkits for any animal species (Figure 2). We demonstrate the versatility and impact of the FAST system by using it to confirm seven receptor-ligand interactions among twelve checkpoint proteins in devils.

**Figure 2.**
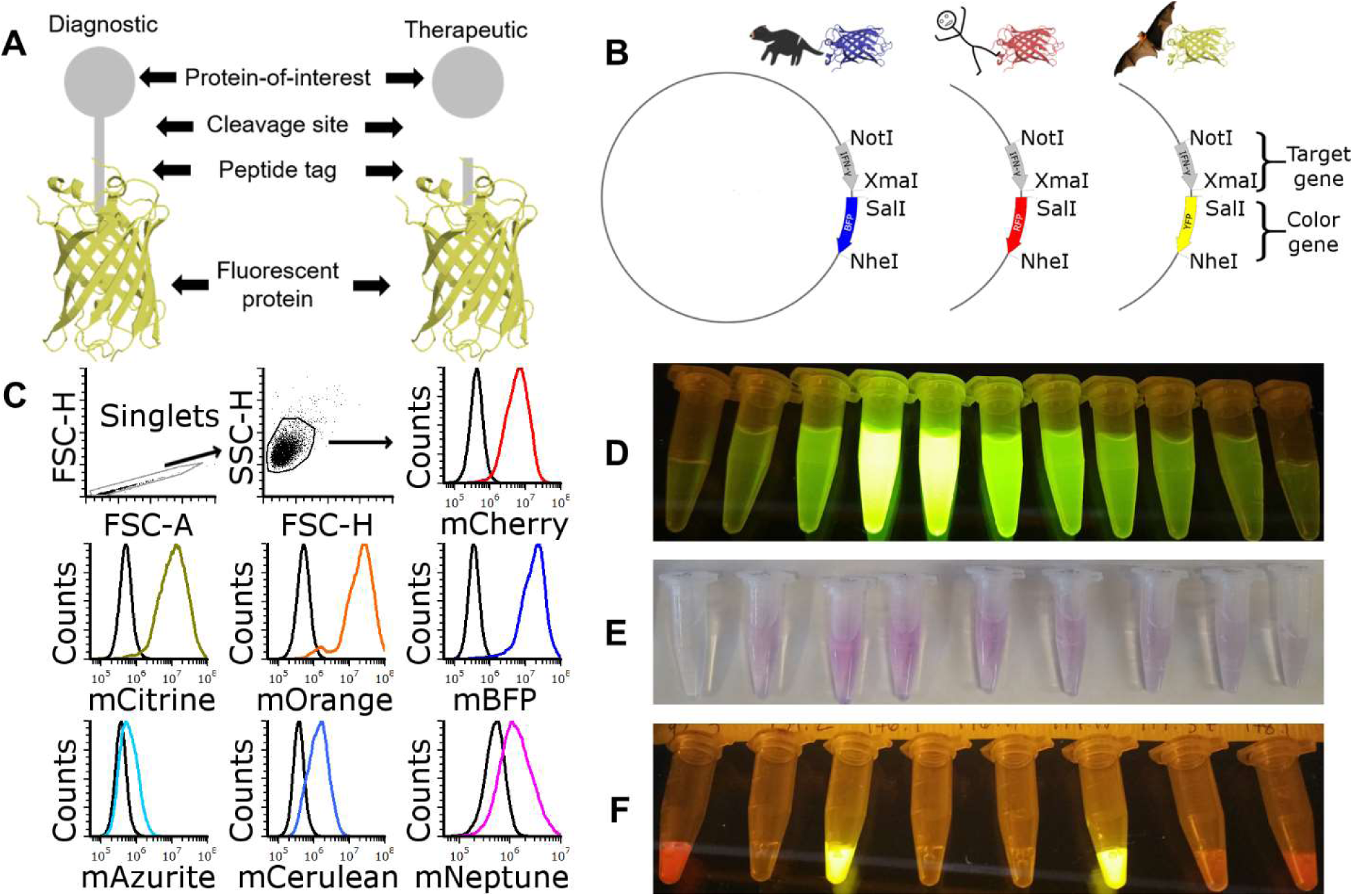
FAST protein schematic and initial testing. (A) Schematic diagram of FAST protein therapeutic and diagnostic (i.e. theranostic) features and (B) vector map showing restriction sites for swapping the target gene (i.e. gene-of-interest) and color genes (i.e. fluorescent protein). (C) Results of flow cytometry binding assay with devil 41BB FAST proteins. The colored lines in the histograms show binding of devil 41BB fused to mCherry, mCitrine, mOrange, mTagBFP, mAzurite, mCerulean3, or mNeptune2 to CHO cells transfected with devil 41BBL, and the black lines show binding to untransfected CHO cells. (D-E) Images showing the gradient of FAST proteins eluted from HisTrap columns. (D) mCitrine excited with blue light and (E) chromogenic visualization of mCherry without excitation. (F) Image of 100 μL of FAST protein excited with blue light (NOTE: mBFP appears clear with blue excitation and amber filter unit).

In humans, these checkpoint proteins have been targets of immunotherapy in clinical trials, but the functional role and binding patterns of these proteins are unknown for most other species. We have used the FAST system to show that the inhibitory checkpoint protein CD200 is highly expressed on DFT cells, opening the door to single-cell phenotyping of circulating tumor cells (CTCs) in devil blood. Furthermore, we are the first to report that co-expression of CD200R1 can block surface expression of CD200 in any species. Understanding how clonal tumor cells graft onto new hosts, evade immune defenses and metastasize within a host will identify evolutionarily conserved immunological mechanisms to help improve cancer, infectious disease, and transplant outcomes for human and veterinary medicine.

## RESULTS

### Fluorescent fusion proteins can be secreted from mammalian cells

Initially, we developed FAST proteins to determine whether monomeric fluorescent proteins could be fused to devil proteins and secreted from mammalian cells (Figure 2A and **Table S2**). We used 41BB (TNFRSF9) for proof-of-concept studies by fusing the extracellular domain of devil 41BB checkpoint molecule to monomeric fluorescent proteins (Figure 2A-B and Figure S1). We used wild-type Chinese hamster ovary (CHO) cells and CHO cells transfected with 41BBL (TNFSF9) to confirm specificity of the 41BB FAST proteins and demonstrate that the fluorescent proteins (mAzurite, mCerulean3, mCherry, mCitrine, mOrange, Neptune2, and mTag-BFP (aka mBFP)) remained fluorescent when secreted from mammalian cells (Figure 2C).

We chose mCherry, mCitrine, mOrange, and mBFP for ongoing FAST protein development. Initial batches of FAST proteins were purified using the 6xHis-tag and eluted with imidazole. The gradient of FAST protein in the collection tubes was apparent when excited with blue light and visualized with an amber filter unit (Figure 2D), allowing immediate confirmation that the fluorescent protein DNA coding sequences were in-frame and the proteins were properly folded. mCherry was visible without excitation or filters (Figure 2E). After combining, concentrating, and sterile filtering the eluted fractions, 100 μL was aliquoted and visualized again using blue light to confirm fluorescent signal (Figure 2F). A full step-by-step protocol and set of experimental templates for creating and testing FAST proteins for any species is available online with the supplementary material.

### Receptor-ligand binding confirmed in single-step staining assays

We chose additional immune checkpoint molecules for FAST protein development (Figure 3A) based on targets of clinical trials and sequence analysis of devil genes ^27, 34, 35^. We transfected the FAST protein expression vectors (**Table S2**) into CHO cells and tested the supernatant against CHO cell lines expressing full-length receptors. 41BB FAST proteins in supernatant exhibited strong binding to 41BBL cell lines, but the fluorescent signals from most other FAST proteins were too weak to confirm binding to the expected receptors (Figure S2). As FAST proteins do not require secondary reagents, we next incubated target cells with purified FAST proteins and added chloroquine to block the lysosomal protein degradation pathway ^36^. This allowed us to take advantage of receptor-mediated endocytosis, which can allow accumulation of captured fluorescent signals inside the target cells ^37^. This protocol adjustment allowed confirmation that CD47-mCherry, CD200-mBFP, CD200-mOrange, CD200R1-mBFP, and CD200R1-mOrange, and PD1-mCitrine bound to their expected receptors (Figure 3B). We also demonstrated the flexibility of the FAST proteins by showing that alternative fusion conformations (Figure S1C-D), such as type II proteins (e.g. mCherry-41BBL) and a devil Fc-tag (e.g. CD80-Fc-mCherry) bound to their expected ligands (Figure 3B). The stability of the fusion proteins was demonstrated using supernatants that were stored at 4 °C for two months prior to use in a one-hour live-culture assay with chloroquine (Figure S3).

**Figure 3.**
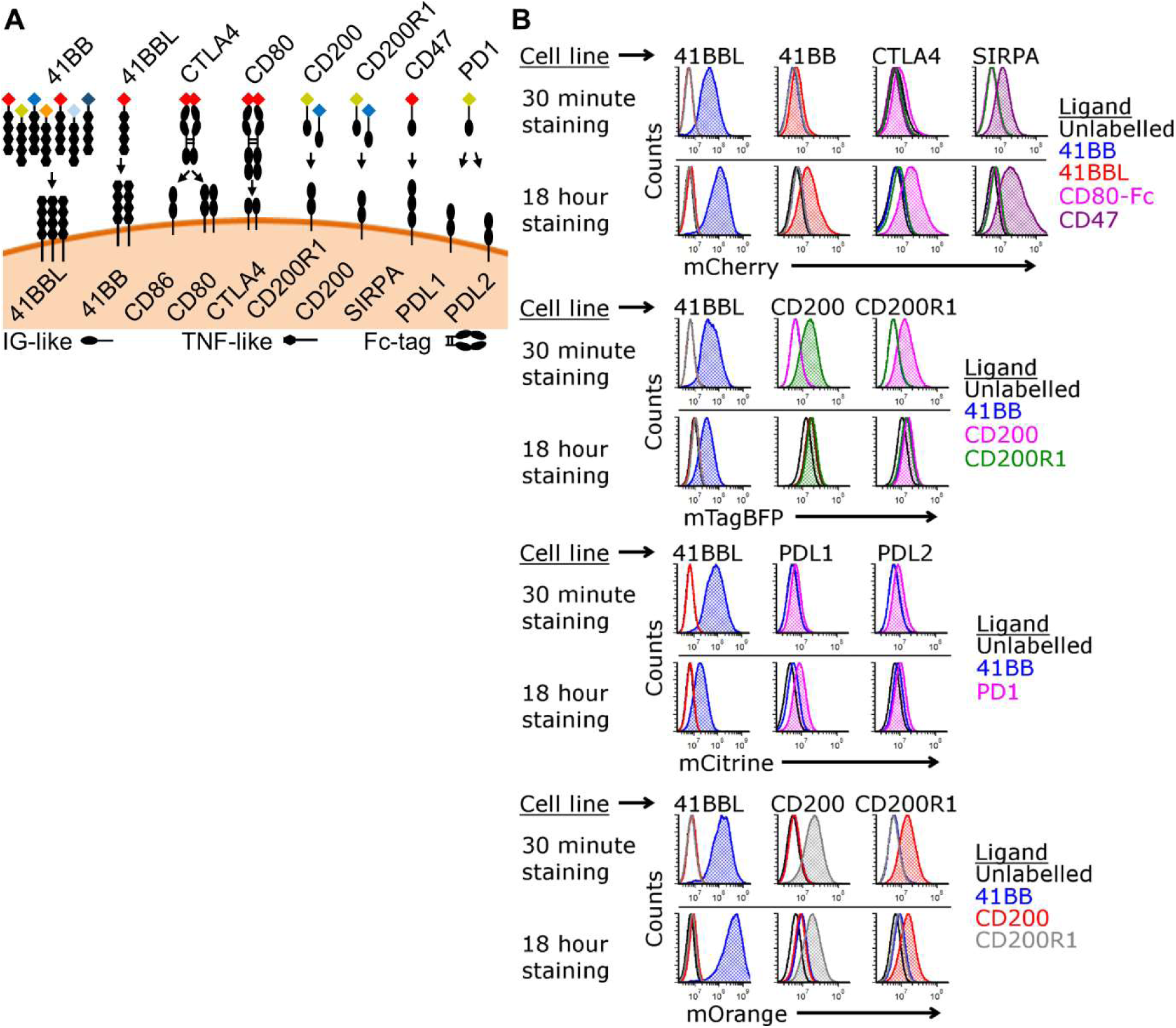
Map and testing of soluble proteins FAST proteins. (A) Diagram of soluble FAST proteins and full-length proteins used for testing of FAST proteins. 41BBL is a type II transmembrane protein; all other proteins are type I. CD80 and CTLA4 soluble FAST proteins included a devil IgG Fc-tag. Arrows indicate interactions confirmed in this study. (B) Histograms showing binding of FAST proteins to CHO cells expressing full-length devil proteins. Target CHO cells were cultured with chloroquine to block lysosomal degradation of FAST proteins and maintain fluorescent signal during live-culture binding assays with 2 μg/well of purified FAST proteins for 30 minutes or 18 hours to assess receptor-ligand binding (n=1/time point).

### Cell lines secreting FAST proteins confirm protein interactions in live coculture assays

To further streamline the reagent development process, we next took advantage of the single-step nature of FAST proteins (i.e. no secondary antibodies or labels needed) in live-cell coculture assays (Figure 4A). Cell lines secreting 41BB-mCherry, 41BBL-mCherry, or CD80-Fc-mCherry FAST proteins were mixed with cell lines expressing full-length 41BB, 41BBL, or CTLA4-mCitrine and cocultured at a 1:1 ratio overnight with chloroquine. Singlet cells were gated (Figure 4B) and binding of mCherry FAST proteins to CFSE or mCitrine-labelled target cells was analyzed (Figure 4C). The strongest fluorescent signal from 41BB-mCherry, 41BBL-mCherry and CD80-Fc-mCherry were detected when cocultured with their predicted receptors, 41BBL, 41BB and CTLA4, respectively.

**Figure 4.**
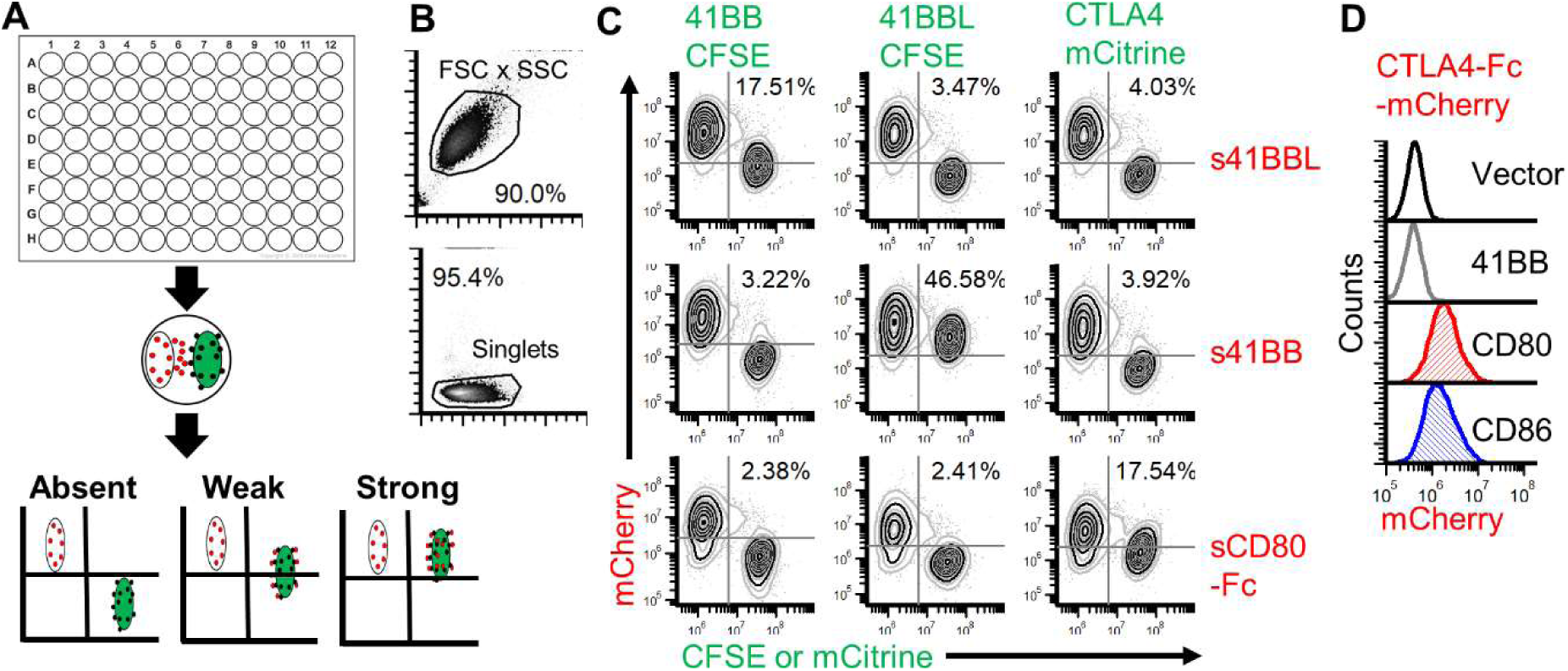
Live-cell coculture assays with FAST proteins. (A) Schematic of coculture assays to assess checkpoint molecule interactions (Absent, Weak, Strong). Cells were mixed and cultured overnight with chloroquine. Protein binding and/or transfer were assessed using flow cytometry. (B) Gating strategy for coculture assays. (C) CHO cells that secrete 41BBL-mCherry, 41BB-mCherry, or CD80-Fc-mCherry were cocultured overnight with target CHO cells that express full-length 41BB, 41BBL, or CTLA4. 41BB and 41BB-L were labeled with CFSE, whereas full-length CTLA4 was directly fused to mCitrine. Cells that secrete mCherry FAST proteins appear in the upper left quadrant. Cells expressing full-length proteins and labeled with CFSE or mCitrine appear in the lower right quadrant. Cells in the upper right quadrant represent binding of mCherry FAST proteins to full-length proteins on CFSE or mCitrine labeled cells. Results shown are representative of n=3/treatment. (D) CTLA4-Fc-mCherry FAST protein binding to DFT cells. DFT1 C5065 cells transfected with control vector (black), 41BB (gray), CD80 (red), or CD86 (blue) were stained with CTLA4-Fc-mCherry supernatant with chloroquine. Results are representative of n=2 replicates/treatment.

### Optimization of the FAST-Fc construct

The fluorescent binding signal of CD80-Fc-mCherry was lower than expected, so we next re-examined our Fc-tag construct. In humans and all other mammals examined to date the IgG heavy chain has glycine-lysine (Gly-Lys) residues at the C-terminus ^38^; the initial devil IgG constant region sequence available to us had an incomplete C-terminus, and thus our initial CD80-Fc-mCherry vector did not have the C-terminal Gly-Lys. We subsequently made a new FAST-Fc construct with CTLA4-Fc-mCherry, which exhibited strong binding to both CD80 and CD86 transfected DFT cells (Figure 4D).

### CD200 mRNA and protein are highly expressed in DFT cells

Analysis of previously published devil and DFT cell transcriptomes suggested that CD200 mRNA is highly expressed in DFT2 cells and peripheral nerves, moderately expressed in DFT1 cells, and lower in other healthy devil tissues (Figure 5A) ^34, 35, 39^. As CD200 is an inhibitory molecule expressed on most human neuroendocrine neoplasms ^40^, and both DFT1 and DFT2 originated from Schwann cells ^35, 41^, we sought to investigate CD200 expression on DFT cells at the protein level. Staining of wild-type DFT1 and DFT2 cells with CD200R1-mOrange FAST protein showed minimal fluorescent signal (Figure 5B). However, overexpression of CD200 using a human EF1α promoter yielded a detectable signal with CD200R1-mOrange binding to CD200 on DFT1 cells. A weak signal from CD200-mOrange was detected on DFT1 cells overexpressing CD200R1 (Figure 5B). To confirm naturally-expressed CD200 on DFT cells we digested CD200 and 41BB FAST proteins using TEV protease to remove the linker and fluorescent reporter. The digested proteins were then used to immunize mice for polyclonal sera production. We stained target CHO cell lines with pre-immune (PI) or immune (I) mouse sera collected after 3X immunizations. Only the immune sera showed strong binding to the respective CD200 and 41BB target cell lines (Figure 5C). After the final immunization (4X), we collected another batch of sera and tested it on DFT1 and DFT2 cells (Figure 5D). In agreement with the transcriptomic data for DFT cells ^34^, the polyclonal sera revealed high levels of CD200 on DFT cells, but low levels of 41BB.

**Figure 5.**
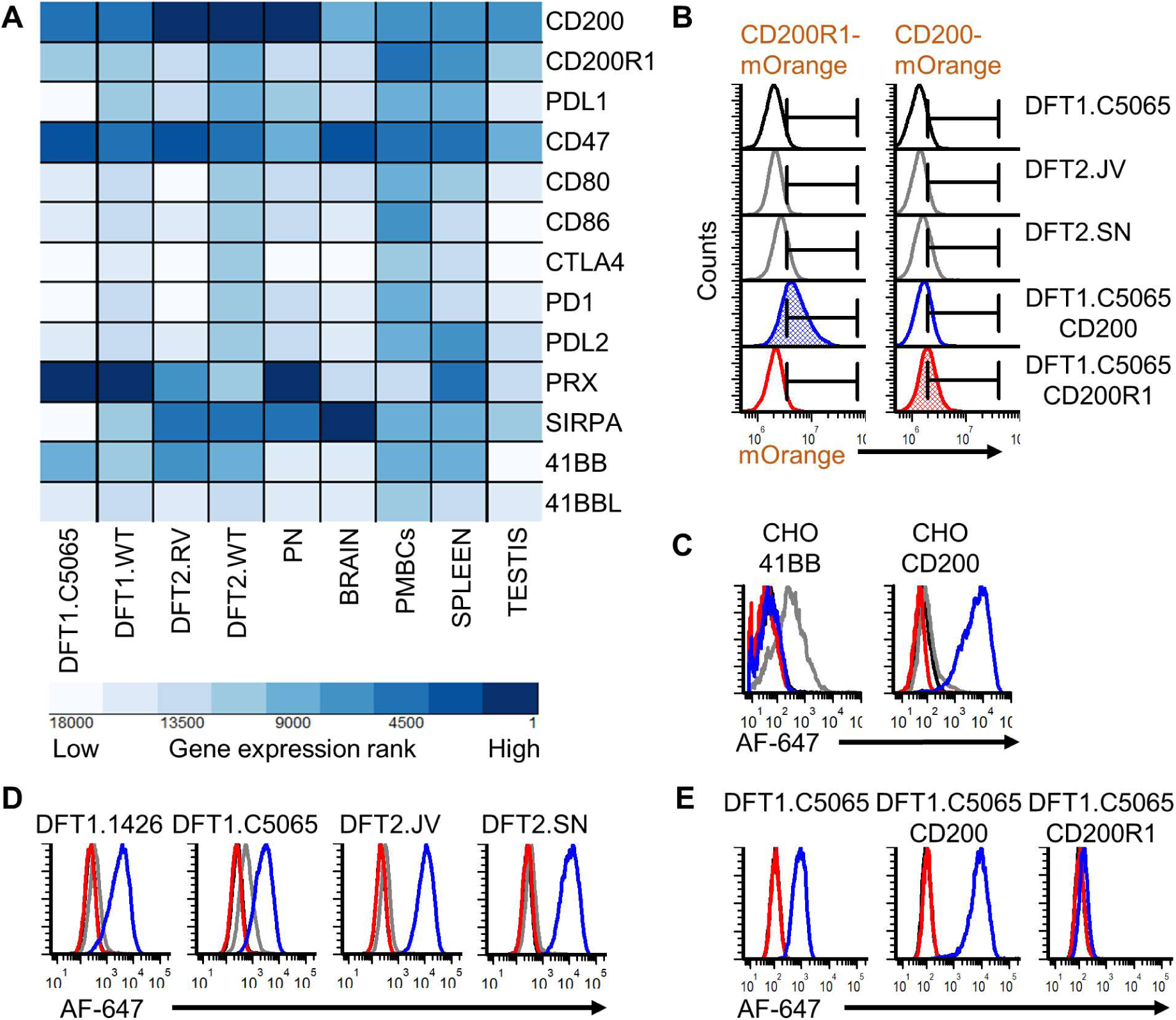
Elevated CD200 expression on DFT cells. (A) Heatmap showing within sample transcript ranking (1 = highest expression) according to RPKM-normalized mRNA sequencing counts of 18,788 annotated coding genes (devil_refv7.0; GCA_000189315.1). Genes-of-interest for this study are plotted as heatmap with dark blue indicating the most highly expressed genes. Technical replicates (n=2) were used for all tissues, except peripheral nerve (PN) (n=1). (B) Wild type DFT1.C5065, DFT2.JV, DFT2.SN, and DFT1.C5065 transfected to overexpress CD200 or CD200R1 were stained with 5 μg of either CD200R1-mOrange or CD200-mOrange FAST protein. Histograms filled with blue or red highlight the cells overexpressing CD200 or CD200R1 and their expected binding interactions with CD200R1 and CD200, respectively. Target cells were cultured with FAST proteins in chloroquine, incubated 30 minutes, and run without washing. Results are representative of n=2 replicates/treatment. (C) Mice were immunized with TEV digested 41BB or CD200 FAST proteins. Black = pre-immune (PI) and gray = immune (I) sera from a mouse immunized with 41BB; red = pre-immune (PI) and blue = immune (I) sera from a mouse immunized with CD200. CHO cells transfected with either full-length 41BB or CD200 were stained with sera and then anti-mouse AlexaFluor-647. Results are representative of n=2/treatment. (D) Sera was used to screen two strains of DFT1 and two strains of DFT2 cells for 41BB and CD200 expression. PI sera was negative in all cases, whereas all DFT1 and DFT2 cells expressed CD200. Results are representative of n=3/treatment. (E) DFT1 C5065 transfected with either vector control, CD200, or CD200R1 were stained with purified polyclonal anti-CD200 and anti-mouse IgG AlexaFluor 647 (black = no antibodies, red = secondary antibody only, blue = primary + secondary antibody). Results are representative of n=2/treatment.

### Overexpression of CD200R1 blocks surface expression of CD200

In humans, overexpression of some checkpoint proteins can block surface expression of heterophilic binding partners in *cis* (e.g. CD80 and PDL1) ^42, 43^. As a potential route for disrupting the inhibitory effects of CD200 on anti-tumor immunity, we tested if overexpression of CD200R1 on DFT cells could reduce CD200 surface expression. We stained a DFT1 strain C5065, and DFT1 C5065 cells transfected to overexpress CD200 or CD200R1 with polyclonal anti-CD200 sera and secondary anti-mouse IgG AF647. We detected no surface protein expression of CD200 DFT1 cells overexpressing CD200R1 (Figure 5E).

### Identification of DFT cells in whole blood using anti-CD200

In addition to high expression of CD200 on neuroendocrine neoplasms ^40, 44^, CD200 is used as a diagnostic marker for several human blood cancers ^45–47^. DFT cells metastasize in the majority of cases ^48^ and our transcriptome results (Figure 5A) suggest that CD200 mRNA is more highly expressed in DFT cells than in peripheral blood mononuclear cells (PBMCs) ^34, 35^. As a result, we tested if CD200 could be used to identify DFT cells in blood. We stained PBMCs and DFT2 cells separately with polyclonal anti-CD200 sera and anti-mouse AlexaFluor 647 and then analyzed CD200 expression by flow cytometry (Figure S4A). We then mixed the stained PBMCs and DFT2 cells at ratios of 1:10 (Figure S4A) and 1:5 (Figure S4B) and analyzed the mixed populations. PBMCs showed minimal CD200 expression and background staining (Figure S4), whereas CD200 was highly expressed on DFT2 cells. CD200+ DFT2 cells were readily distinguishable from PBMCs.

As our RNAseq results only included mononuclear cells, we next performed a pilot test to determine if DFT cells could be spiked into whole devil blood and identified via flow cytometry using CD200 staining. DFT1 and DFT2 cells were labeled with CellTrace violet (CTV) and 10,000 cells were diluted directly into 100 μL of whole blood from a healthy devil (n=1/treatment; n = 1 devil). The cells were then stained with purified polyclonal anti-CD200 with and without secondary anti-mouse IgG AF647 prior to red blood cell (RBC) lysis. Initial results showed that DFT2 cells expressed CD200 above the leukocyte background, but that DFT1 cells could not be distinguished from leukocytes (Figure S5). To eliminate the secondary antibody step from the whole blood staining protocol we next labelled the polyclonal anti-CD200 and normal mouse serum (NMS) with a no-wash Zenon mouse IgG AF647 labeling reagent (n = 1/treatment; n = 2 devils). This system again showed that CD200 expression could be used to identify DFT2 cells in blood (Figure 6), suggesting that CD200 is a candidate marker for identification of metastasizing DFT2 cells.

**Figure 6.**
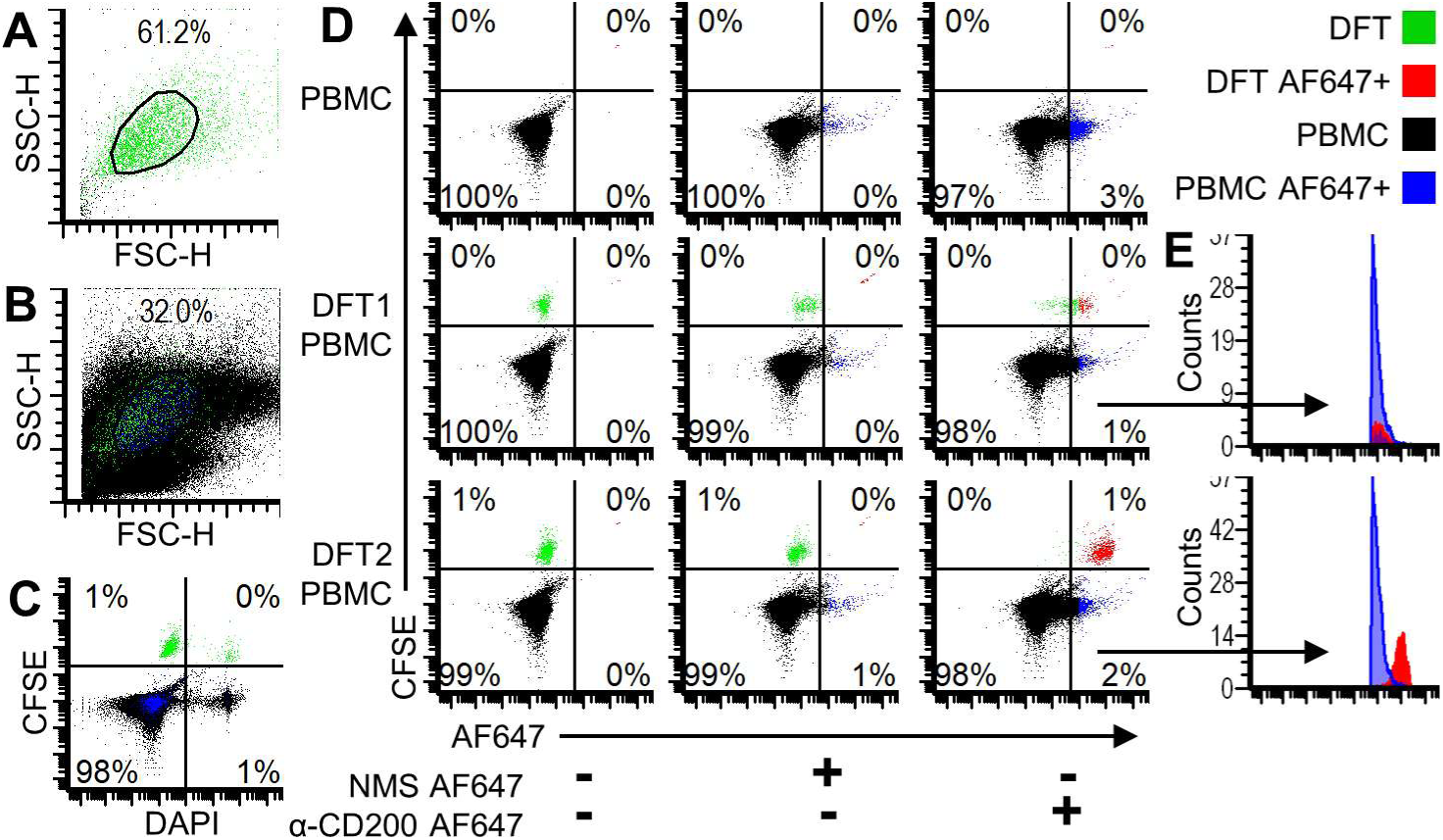
CD200 identifies DFT cells in whole blood. Color dot plots showing DFT cells in green (CFSE), PBMCs in black, DFT Alexa Fluor 647+ (AF647) cells in red, and PBMC AF647+ in blue. (A) Forward-and side-scatter plot of DFT.JV cells and (B) DFT.JV cells mixed with PBMCs. (B) Color dot plot showing dead cells stained with DAPI (right quadrants) and CFSE-labeled DFT cells (upper quadrants). (D) The top row shows unmixed PBMCs. The middle row and bottom row show DFT1.C5065 and DFT2.JV cells, respectively, mixed with PBMCs. Alexa Fluor 647+ DFT (red) and PBMC (blue) are in the right quadrants. (E) Histogram overlays to highlight AF647+ (right quadrants) from DFT1-PBMC and DFT2-PBMC mixtures. Cells were analyzed on the Beckman-Coulter MoFlo Astrios.

## DISCUSSION

Naturally occurring cancers provide a unique opportunity to study immune evasion and the metastatic process across diverse hosts and environments. The exceptionally high cancer rate in Tasmanian devils coupled with the two transmissible tumors currently circulating in the wild warrants a thorough investigation of the devil immune system. However, taking advantage of such natural disease models has been out of reach for most species due to a lack of reagents. The FAST protein system we developed here is well-suited to discovering additional DFT markers, and more generally, filling the reagent gap for non-traditional species. For proteins like 41BB that have high affinity for 41BBL, FAST proteins can be used as detection reagents directly from supernatant. For other molecules with lower receptor-ligand affinity, the FAST proteins can be purified, digested with a protease to remove the non-target proteins and used for production of polyclonal or monoclonal antibodies.

The simple cut-and-paste methods for vector assembly lend the FAST protein system to entry level immunology and molecular biology skill sets. Additionally, the ability of FAST proteins to be used in live coculture assays and with elimination of secondary reagents, will increase efficiency and reduce experimental error for advanced human and mouse cancer immunology studies. For example, previous high-throughput studies have used a two-step staining process (i.e. recombinant protein + secondary antibody) to screen more than 2000 protein interactions ^49, 50^; this type of assay can be streamlined using FAST proteins to eliminate the need for secondary antibodies. Fc-tags or other homodimerization domains can be incorporated into FAST proteins to increase binding for low-affinity interactions.

Production of recombinant proteins in cell lines that closely resemble the physiological conditions of the native cell type (i.e. mammalian proteins produced in mammalian cell lines) are more likely to yield correct protein folding, glycosylation, and function than proteins produced using evolutionarily distant cell lines ^51^. The fluorescent fusion proteins developed here that take advantage of natural receptor expression and cycling processes (e.g. CTLA4 transendocytosis) in eukaryotic target cells; bacterial protein production methods are not amenable to coculture with eukaryotic target cells in immunological assays ^52^. Our demonstration of the FAST protein system in CHO cells, which are used to produce approximately 70% of recombinant pharmaceutical proteins ^53^, suggest that this method can be efficiently integrated into existing research and development pipelines for humans and other vertebrate species.

A primary question in transmissible tumor research is why genetically mismatched cells are not rejected by the host. Successful infection of devils with DFT cells relies on the ability of the tumor allograft to evade and manipulate host defences. The “missing-self” hypothesis suggests that the lack of constitutive MHC-I expression on DFT1 cells should lead to natural killer (NK) cell-mediated killing of the allograft tumor cells. Here we used the FAST protein system to develop a tool set to address this question and show that DFT1 and DFT2 cells express CD200 at higher levels than most other devil tissues examined to date. CD200 has been shown to directly inhibit NK cells in other species ^54–56^, so overexpression of CD200 is a potential mechanism of immune evasion of NK responses by DFT cells.

We hypothesize that CD200 could be particularly important in DFT transmission as the CD200-CD200R pathway is critical to the initial stages of establishing transplant and allograft tolerance in other species ^57–59^. In line with this hypothesis, a recent study reported that overexpressing several checkpoint molecules, including CD200, PDL1, and CD47, in mouse embryonic stem cells could be used to generate teratomas that could establish long-term allograft tolerance in fully immunocompetent hosts ^60^. We have previously reported that PDL1 mRNA and protein are upregulated on DFT2 cells in response to IFNγ ^25^, and our transcriptome results show that CD47 is expressed at moderate to high levels in DFT cells. Here we show that overexpression of CD200R1 on DFT1 eliminates binding of our polyclonal anti-CD200 antibodies, suggesting that DFT cells overexpressing CD200R1 could be used to test the role of CD200 in allograft tolerance. Alternatively, genetic ablation of CD200 in DFT cells could be used as a complementary approach to examine the role of immune checkpoint molecules in DFT allograft tolerance. The CD200-CD200R1 pathway has been implicated in reducing IFNγ production by dendritic cells ^59^ and decreasing the responsiveness of myeloid cells to IFNγ stimulation ^61^. Low MHC-I expression is a primary means of immune evasion by DFT1 cells, and disrupting the CD200-CD200R1 pathway could facilitate improved recognition of DFT1 cells by CD8 T cells by enhancing IFNγ-mediated MHC-I upregulation. Recent work in mice has identified immunosuppressive natural regulatory plasma cells that express CD200, LAG3, PDL1, and PDL2; we have previously identified PDL1+ cells with plasma cell morphology near or within the DFT microenvironment ^25^.

Previous DFT vaccine efforts have used killed DFT cells with adjuvants ^62, 63^. A similar approach to treat gliomas in dogs reported that tumor-lysate with CD200 peptides nearly doubled progression-free survival compared to tumor lysate alone ^64^. Like devils, several breeds of dog are prone to cancer and these genetically-outbred large animal models provide a fertile ground for testing cancer therapies. Interestingly, the CD200 peptides are reported to provide agonistic function through CD200-like activation receptors (CD200R4) rather than by blocking CD200R1 ^44, 64^. The functional role of CD200-CD200R pathway in devils remains to be elucidated, but the CD200R1_NPLY_ inhibitory motif and key tyrosine residues are conserved in devil CD200R ^27, 65, 66^, demonstrating this motif is conserved over 160 million years of evolutionary history ^67^. In addition to agonistic peptides, several other options for countering CD200-CD200R immune inhibition are possible. Human chronic lymphocytic leukemia cells often express high levels of CD200, which can be downregulated in response to imiquimod ^68^. Likewise, we have previously shown that DFT1 cells downregulate expression of CD200 mRNA *in vitro* in response to imiquimod treatment ^34^.

In mice, chronic salmonella and schistosoma infections upregulated both CD200 and CD200R ^69^. Several viruses, including cytomegalovirus ^70^ and herpesvirus ^71^ manipulated the CD200-CD200R pathway as a means of immune evasion. Interestingly, in one of the longest running and most in-depth studies of host-pathogen coevolution, CD200R was shown to be under selection in rabbits in response to myxoma virus biocontrol agent ^72^. As DFT1 and DFT2 have been circulating in devils for more than 20 years and 5 years, respectively, it will be important to monitor CD200/R expression and the potential evolution of paired activating and inhibitory receptors in these natural disease models ^73^.

Immunophenotyping and single-cell RNAseq of circulating tumor cells (CTCs) has potential to identify key gene expression patterns associated with metastasis and tissue invasion. Periaxin (PRX) is the most sensitive and specific marker for DFT1 cells in immunohistochemistry assays ^74^. Unfortunately, PRX is expressed primarily in the cytoplasm, which eliminates the possibility of using PRX as a marker to sort live cells via flow cytometry for single-cell RNAseq. However, CD200 is a potential marker for the identification of circulating tumor cells (CTCs) from devil blood. As proof of concept, DFT2 cells could be identified in devil blood spiked with DFT2 cells. As CTCs are likely to be rare in the blood of most infected devils, CD200 alone would be insufficient for identifying DFT1 cells. Additional surface DFT markers would be required to purify CTCs for metastases and tissue invasion analyses. The FAST protein system provides a simple procedure to facilitate the production of a panel of DFT-markers to help identify key proteins in the metastatic process.

In summary, the simple “cut-and-paste” production of the vectors and single-step testing pipeline of the FAST system provided multiple benefits. The FAST system allowed us to characterize receptor-ligand interactions, and to identify evolutionarily conserved immune evasion pathways in naturally occurring transmissible cancers. Our initial implementation of the system confirmed numerous predicted protein interactions for the first time in a marsupial species and documented high expression of the inhibitory molecule CD200 on DFT cells. The high expression of CD200 in devil nervous tissues and neuroendocrine tumors, downregulation of CD200 in response to imiquimod, and binding of CD200 to CD200R1, is consistent with results from human and mouse studies. Consequently, the CD200/R pathway provides a promising immunotherapy and vaccine target for DFTs. Beyond this study, FAST proteins meet the key attributes needed for reagent development, such as being straightforward to make, stable, versatile, renewable, cheap, and amenable to high-throughput testing ^75^. The direct fusion of the reporter protein to the protein-of-interest allows for immediate feedback during transfection, supernatant testing, and protein purification; proteins with frameshifts, introduced stop codons, or folded improperly will not fluoresce and can be discarded after a simple visualization, rather than only after extensive downstream testing. Efficient mapping of immune checkpoint interactions across species can identify evolutionarily conserved immune evasion pathways and appropriate large animal models with naturally occurring cancer. This knowledge could inform veterinary and human medicine in the fields of immunological tolerance to tissue transplants, infectious disease, and cancer.

## MATERIALS AND METHODS

### Study design

The objectives of this study were to fill a major gap in our understanding of the mammalian immune system and to understand how genetically mismatched transmissible tumors evade host immunity. To achieve this goal, we developed a recombinant protein system that directly fuses proteins-of-interest to a fluorescent reporter protein. The first phase was to determine if the fluorescent protein remained fluorescent after secretion from mammalian cells and to confirm that proteins bound to their predicted receptors (i.e. ligands). Initial testing was performed in CHO cells and follow-up assays used devil cells. To further demonstrate the functionality of this system for antibody development, mice were immunized with either 41BB or CD200 proteins. Pre- and post-immunization polyclonal sera was used to confirm that the proteins used for immunization induced antibodies that specifically bound to surface-expressed recombinant proteins and native proteins on devil facial tumor cells.

### Target transcript amplification

Target gene DNA sequences for vector construction were retrieved from Genbank, Ensembl or de novo transcriptome assemblies (**Table S2**) ^76^. Target DNA was amplified from a cDNA template or existing plasmids using primers and PCR conditions shown in **Tables S2-S4** using Q5 High-Fidelity 2X Master Mix (New England Biolabs # M0494L). Primers were ordered with 5’ base extensions that overlapped expression vectors on either side of the restriction sites. The amplified products were identified by gel electrophoresis and purified using Nucleospin PCR and Gel Clean Up Kit (Macherey-Nagel # 740609.5). Alternatively, DNA sequences were purchased as double stranded DNA gblocks (**Table S5**) (Integrated DNA Technologies) for direct assembly into expression vectors.

### Construction of all-in-one Sleeping Beauty transposon vectors

All new plasmids were assembled using NEBuilder kit (NEB # E5520S) following the manufacturer’s recommendations unless otherwise noted. DNA inserts, digested plasmids, and NEBuilder master mix were incubated for 60 minutes at 50 °C and then transformed into DH5α included with the NEBuilder kit. Plasmid digestions were performed following manufacturer recommendations and generally subjected to Antarctic phosphatase (New England Biolabs # M0289S) treatment to prevent potential re-annealing. Sleeping Beauty transposon vectors pSBbi-Hyg (Addgene # 60524), pSBbi-BH (Addgene # 60515), pSBtet-Hyg (Addgene # 60508), pSBtet-RH (Addgene # 60500) were gifts to Addgene from Eric Kowarz ^77^. The pCMV(CAT)T7-SB100 containing the CMV promoter and SB100X transposase was a gift to Addgene from Zsuzsanna Izsvak (Addgene # 34879) ^78^. We first constructed an all-in-one Sleeping Beauty vector by inserting a CMV promoter and SB100X transposase from pCMV(CAT)T7-SB100 ^78^ into pSBi-BH ^77^ (**Tables S3-S4**). This was accomplished by using pAF111-vec.FOR and pAF111.1.REV primers to amplify an overlap region from pSBbi-BH (insert 1) and pAF111-2.FOR and pAF111-2.REV to amplify the CMV-SB100X region from pCMV(CAT)T7-SB100 (insert 2). The purified amplicons were then used for NEBuilder assembly of pAF111. The final all-in-one vectors pAF112 (hygromycin resistance and luciferase) and pAF123 (hygromycin resistance) were assembled from the pAF111 components. pAF112 was assembled by amplifying the Luc2 luciferase gene (insert 1) from pSBtet-Hyg and the P2A-hygromycin resistance gene (insert 2) from pSBbi-BH and inserting into the pAF111 Bsu36I digest using NEBuilder. pSBbi-Hyg was Bsu36I-digested to obtain the hygromycin resistance gene, and this fragment was inserted into Bsu36I-digested pAF111 using T4 ligase cloning to replace the BFP-P2A-hygromycin segment in pAF111.

### Construction of full-length protein vectors

All full-length gene coding sequences except CTLA4 were cloned into the a pAF112 SfiI digest (**Table S2**). All full-length vectors also contain luciferase with T2A peptide linked to the hygromycin resistance protein; luciferase was included for use in downstream functional testing that was not part of this study. Tasmanian devil CTLA4 was cloned into a NotI-HF and XmaI digest of pAF100 that was used in a different study but is derived from vectors pAF112 and pAF138. Additionally, we also used devil PDL1 (CHO.pAF48) and 41BBL (CHO.pAF56) cell lines developed using a vector system described previously ^25^.

### Construction of FAST protein vectors

Plasmids containing fluorescent protein coding sequences mCerulean3-N1 (Addgene # 54730), mAzurite-N1 (Addgene # 54617), mOrange-N1 (Addgene # 54499), mNeptune2-N1 (Addgene # 54837) were gifts to Addgene from Michael Davidson. mTag-BFP was amplified from pSBbi-BH, mCitrine was amplified from pAF71, and mCherry was amplified from pTRE-Dual2 (Clontech # PT5038-5). pAF137 was constructed by amplifying the devil 41BB extracellular domain with primers pAF137-1.FOR and pAF137-1.REV and amplifying mCherry with pAF137-2a.FOR and pAF137-2.REV (**Table S3-4**). 5’ extensions on pAF137-1.FOR and pAF137-2.REV were used to create overlaps for NEBuilder assembly of pAF137 from a pAF123 SfiI-digested base vector. 3’ extensions on pAF137-1.REV and pAF137-2a.FOR were used to create the linker that included an XmaI/SmaI restriction site, TEV cleavage tag, GSAGSAAGSGEF linker peptide, and 6x-His tag between the gene-of-interest and fluorescent reporter. The GSAGSAAGSGEF was chosen due to the low number of large hydrophobic residues and less repeated nucleic acids than are needed with other flexible linkers such as (GGGS)_4_ ^79^. The pAF137 primer extensions also created 5’ NotI and 3’ NheI sites in the FAST vector to facilitate downstream swapping of functional genes and to create a Kozak sequence ^80^ (GCCGCCACC) upstream of the FAST protein open-reading frame. Following confirmation of correct assembly via DNA sequencing, the FAST 41BB-mCherry (pAF137) was digested and used as the base vector (Figure 2B and Figure S1A-B) for development of FAST vectors with alternative fluorescent proteins. This was accomplished by digestion of pAF137 with SalI and NheI and then inserting PCR-amplified coding sequences for other fluorescent proteins using NEBuilder (**Tables S3-S4**).

Type I FAST (extracellular N-terminus, cytoplasmic C-terminus) protein vectors were constructed by digestion of 41BB FAST vectors with NotI and either XmaI or SmaI (Figure 2B and Figure S1A-B), and then inserting genes-of-interest (**Tables S2-4**). To create an Fc-tagged FAST protein we fused the extracellular domain of devil CD80 to the Fc region of the devil IgG (Figure S1C). The Fc region was amplified from a devil IgG plasmid provided by Lynn Corcoran (Walter and Eliza Hall Institute of Medical Research). All secreted FAST proteins in this study used their native signal peptides, except for 41BBL. 41BBL is a type II transmembrane protein in which the signal peptide directly precedes the cytoplasmic and transmembrane domains of the protein (cytoplasmic N-terminus, extracellular C-terminus). As type I FAST vectors cannot accommodate this domain architecture, we developed an alternative base vector for type II transmembrane FAST proteins (Figure S1D). To increase the probability of efficient secretion of type II FAST proteins from CHO cells, we used the hamster IL-2 signal peptide (accession # NM_001281629.1) at the N-terminus of the protein, followed by a SalI restriction site, mCherry, an NheI restriction site, 6x-His tag, GSAGSAAGSGEF linker, TEV cleavage site, XmaI/SmaI restriction site, the gene-of-interest, and a PmeI restriction site following the stop codon.

### General plasmid assembly, transformation, and sequencing

Following transformation of assembled plasmids, colony PCR was performed as an initial test of the candidate plasmids. Single colonies were inoculated directly into a OneTaq Hot Start Quick-Load 2X Master Mix (NEB # M0488) with primers pSB_EF1a_seq.FOR (atcttggttcattctcaagcctcag) and pSB_bGH_seq.REV (aggcacagtcgaggctgat). PCR was performed with 60 °C annealing temperature for 25-35 cycles. Colonies yielding appropriate band sizes were used to inoculate Luria broth with 100 μg/mL ampicillin for bacterial outgrowth overnight at 37 °C and 200 RPM. The plasmids were purified using standard plasmid kits and prepared for Sanger sequencing using BigDye Terminator v3.1 Cycle Sequencing Kit (ThermoFisher # 4337455) with pSB_EF1a_seq.FOR and pSB_bGH_seq.REV primers. The BigDye® Terminator was removed using Agencourt CleanSEQ® (Beckman Coulter # A29151) before loading samples to an Applied Biosystems® 3500XL Genetic Analyzer (Applied Biosystems) for sequencing by fluorescence-based capillary electrophoresis.

### General cell culture conditions

DFT1 cell line C5065 and DFT2 cell line JV were cultured at 35 °C with 5% CO_2_ in cRF10 (10% complete RPMI (Gibco # 11875-093) with 2 mM L-glutamine, supplemented with 10% heat-inactivated fetal bovine serum (FBS), and 1% antibiotic-antimycotic (ThermoFisher # 15240062). RPMI without phenol red (Sigma # R7509) was used to culture FAST protein cell lines when supernatants were collected for downstream flow cytometry assays. Devil peripheral blood cells were cultured in cRF10 at 35 °C with 5% CO_2_. CHO cells were cultured at 37 °C in cRF10 during transfections and drug selection but were otherwise cultured at 35 °C in cRF5 (5% complete RPMI). For production of purified recombinant proteins, stably transfected CHO cells were cultured in suspension in spinner flasks in chemically defined, serum-free CHO Ex-Cell (Sigma # 14361C) media supplemented with 8 mM L-glutamine, 10 mM HEPES, 50 μM 2-ME, 1% (v/v) antibiotic-antimycotic, and 1 mM sodium pyruvate and without hygromycin.

### Transfection and Generation of Recombinant Cell Lines

Stable transfections of CHO and DFT cells were accomplished by adding 3 x 10^5^ cells to each well in 6-well plates in cRF10 and allowing the cells to adhere overnight. The next day, 2 μg of plasmid DNA was added to 100 μL of PBS in microfuge tubes. Polyethylenimine (PEI) (linear, MW 25,000; Polysciences # 23966-2) was diluted to 60 μg/mL in PBS and incubated for at least two minutes. 100 μL of the PEI solution was added to the 100 μL of plasmid DNA in each tube to achieve a 3:1 ratio of PEI:DNA. The solution was mixed by gentle pipetting and incubated at room temperature for 15 minutes. Whilst the solution was incubating, the media on the CHO cells were replaced with fresh cRF10. All 200 μL from each DNA:PEI mix was then added dropwise to the CHO cells and gently rocked side-to-side and front-to-back to evenly spread the solution throughout the well. The plates were then incubated overnight at 37 °C with 5% CO_2_. The next day the plates were inspected for fluorescence and then the media was removed and replaced with cRF10 containing 1 mg/mL hygromycin (Sigma # H0654). The media was replaced with fresh cRF10 1 mg/ml hygromycin every 2-3 days for the next seven days until selection was complete. The cells were then maintained in 0.2 mg/mL hygromycin in cRF5 at 35 °C with 5% CO_2_. Supernatant was collected 2-3 weeks post-transfection and stored at 4 °C for two months to assess stability of secreted FAST proteins.

### Protein production and purification

16 days post-transfection the first batch of FAST protein cell lines were adapted to a 1:1 mix of cRF5 and chemically defined, serum free CHO Ex-cell media for 1-2 days to facilitate adaptation of the adherent CHO cells to suspension culture in serum-free media. At least 5×10^7^ cells were then transferred to Proculture spinner flasks (Sigma # CLS45001L, CLS4500250) and stirred at 75 RPM at 35 °C in 5% CO_2_ on magnetic stirring platforms (Integra Bioscience # 183001). Cells were maintained at a density ranging from 5×10^5^ to 2×10^6^ cells/ml for 8-14 days. Supernatant was collected every 2-3 days, centrifuged at 3200 RCF for 10 minutes, stored at 4 °C, and then purified using the ÄKTA start protein purification system (GE Life Sciences # 29022094). The supernatant was diluted 1:1 with 20 mM sodium phosphate pH 7.4 and then purified using HisTrap Excel columns (GE Life Sciences # 17-3712-05) according to the manufacturer’s instructions. Samples were passed through the columns using a flow rate of 2 mL/minute at 4 °C; all wash and elution steps were done at 1 mL/minute. Elution from HisTrap columns (GE Life Sciences # 17-3712-05) was accomplished using 0.5 M imidazole and fractionated into 1 ml aliquots using the Frac30 fraction collector (GE Life Sciences # 29023051). Fluorescence of FAST proteins was checked via brief excitation (Figure 2D) on a blue light transilluminator with an amber filter unit. In the case of mCherry chromogenic color was visible (Figure 2E) without excitation. Fractions containing target proteins were combined and diluted to 15 mL with cold PBS, dialyzed (Sigma # PURX60005) in PBS at 4 °C, 0.22 μm sterile-filtered (Millipore # SLGV033RS) and concentrated using Amicon Ultra centrifugal filter units (Sigma # Z706345). The protein concentration was quantified using the 280 nm absorbance on a Nanodrop spectrophotometer. Extinction coefficients using for each protein were calculated using the ProtParam algorithm ^81^. The proteins were then aliquoted into microfuge tubes and frozen at −80°C until further use. The CTLA4-Fc-mCherry protein was designed, assembled, and tested separately from the other FAST proteins and was tested directly in supernatant without purification.

### Preparation of CHO cells expressing full-length proteins for flow cytometry

CHO cells expressing full-length proteins were thawed in cRF10 and then maintained in cRF5 with 0.2 mg/mL hygromycin. The adherent CHO cells were washed with PBS and incubated with trypsin for 5 minutes at 37 °C to remove cells from the culture flask. Trypsin was diluted 5X with cRF5 and centrifuged at 200 RCF for 5 minutes. Cells were resuspended in cRF5, counted (viability > 95% in all cases), and resuspended and aliquoted for assays as described below.

### Initial staining of CHO cells with 41BB FAST protein supernatants (without chloroquine)

Supernatants (cRF5) were collected from CHO cells expressing devil 41BB-extracellular domain fused to either mCherry (pAF137), mCitrine (pAF138), mOrange (pAF164), mBFP (pAF139), mAzurite (pAF160), mCerulean3 (pAF161), or mNeptune2 (pAF163) (**Tables S2-4)**. The supernatant was spun for 10 minutes at 3200 RCF to remove cells and cellular debris, and then stored at 4 °C until further use. CHO cells expressing devil 41BBL (CHO.pAF56) and untransfected CHO cells were prepared as described above. Flow cytometry tubes were loaded with 5 x 10^4^ target CHO cells/well in cRF5, centrifuged 500 RCF for 3 minutes, and then resuspended in 200 μL of supernatant from the 41BB FAST cell lines (n=1/treatment). The tubes were then incubated for 15 minutes at 4 °C, centrifuged at 500 RCF for 3 minutes, resuspended in 400 μL of cold FACS buffer, and stored on ice until the data were acquired on a Beckman-Coulter Astrios flow cytometer (Figure 2C). All flow cytometry data was analyzed using FCS Express 6 Flow Cytometry Software version 6 (Denovo Software).

### Staining CHO cells with FAST protein supernatants (without chloroquine)

U-bottom 96-well plates were loaded with 1 x 10^5^ target CHO cells/well in cRF5, centrifuged 500 RCF for 3 minutes, and then resuspended in 175 μL of cRF5 supernatant from FAST cell lines collected 11 days after transfection (n=1/treatment). The plates were then incubated for 30 minutes at room temperature, centrifuged at 500 RCF for 3 minutes, resuspended in 200 μL of cold FACS buffer, centrifuged again and fixed with FACS fix buffer (PBS, 0.02% NaN3, 0.4% formalin, 10g/L glucose). The cells were transferred to tubes, diluted with FACS buffer and analyzed on a Beckman-Coulter Astrios flow cytometer (Figure S2).

### Staining CHO cells with purified FAST proteins (with chloroquine)

Purified FAST proteins were diluted to 20 μg/mL in cRF5, aliquoted into V-bottom 96-well transfer plates, and then stored at 37 °C until target cells were ready for staining. Target cells were resuspended in cRF5 with 100 μM chloroquine and 100,000 cells/well were aliquoted into U-bottom 96-well plates. 100 μL of the diluted FAST proteins (n=1/treatment, 2 timepoints/treatment) were then transferred from the V-bottom plates into the U-bottom 96-well plates containing target cells. The final volumes and concentrations were 200 μL/well in cRF5 with 50 μM chloroquine and 2 μg/well of FAST proteins. One set of plates was incubated at 37 °C for 30 minutes and another set of plates was incubated at 37 °C overnight. The cells were then centrifuged 500 RCF for 3 minutes, the media decanted, and incubated for 5 minutes with 100 μL of trypsin to dislodge adherent cells. The cells were then washed with 200 μL of cold FACS buffer, fixed, resuspended in cold FACS buffer, and transferred to tubes for analysis on the Astrios flow cytometer (Figure 3B).

### Staining CHO cells with FAST supernatants (with chloroquine)

The protocol for using FAST protein supernatants was the same above as the preceding experiment except for the modifications described here. Supernatants were collected 2-3 weeks post-transfection, centrifuged at 3200 RCF for 10 minutes, and stored at 4 °C for 2 months. Prior to staining for flow cytometry, the supernatant was 0.22 μm filtered. Supernatant was then loaded into V-bottom 96-well plates to facilitate rapid transfer to staining plates and stored at 37 °C until target cells were ready for staining. Target cells were prepared as described above except for being diluted in cRF5 with 100 μM chloroquine. 2 x 10^5^ cells/well (100 μL) were then loaded into U-bottom 96-well plates. 100 μL of FAST protein supernatant (n=1/treatment) was then transferred from the V-bottom plates to achieve 50 μM chloroquine and the cells were then incubated at 37°C for 60 minutes. The plates were then washed, fixed, and analyzed on the Astrios flow cytometer (Figure S3). A similar procedure was used for staining stably-transfected DFT cells with CTLA4-Fc-mCherry, except that the supernatant was used fresh (Figure 4D).

### Coculture assay with full-length target and FAST protein CHO cell lines (with chloroquine)

CHO cells expressing full-length CTLA4 with a C-terminal mCitrine, and CHO cells expressing full-length 41BB or 41BBL were labelled with 5 μM CFSE; CFSE and mCitrine were analyzed using the same excitation laser (488 nm) and emission filters (513/26 nm). 1 x 10^5^ FAST protein-secreting cells were mixed with 1 x 10^5^ target cells in cRF5 with 50 μM chloroquine and incubated overnight at 37 °C in 96-well U-bottom plates (Figure 4A). The next day the cells were rinsed with PBS, trypsinized, washed, fixed, and resuspended in FACS buffer prior to running flow cytometry. Cells were gated on forward and side scatter (FSC x SSC) and for singlets (FSC-H x FSC-A). (Figure 4B). Data shown in Figure 4C is representative of n=3 technical replicates/treatment. Data was collected using a Beckman Coulter MoFlo Astrios and analyzed using FCS Express.

### Analysis of checkpoint molecule expression in DFT cells and Tasmanian devil tissues

RNAseq data was generated during previous experiments, aligned against the reference Tasmanian devil genome Devil_ref v7.0 (GCA_000189315.1) and summarised into normalized read counts as previously described ^34, 35^. RPKM-normalized read counts were produced in R using edgeR^82^. Genes were ranked from highest RPKM-normalized count to lowest RPKM-normalized count, and a heat map was produced for the genes of interest using the heatmap.2 function of gplots. Heatmap color represents gene ranking among 18,788 predicted protein-coding genes in the reference genome.

### Staining DFTs cell with CD200/R FAST proteins

50,000 DFT cells/well were aliquoted into u-bottom 96-well plates, washed with 150 μL of cRF10, and resuspended in 100 μL of warm cRF10 containing 100 μM chloroquine. 5 μg of FAST protein/well was then added and mixed by pipetting. The plates were then incubated at 37 °C for 30 minutes. The cells were then transferred to microfuge tubes without washing, stored on ice, and analyzed on a Beckman Coulter MoFlo Astrios (n=2/treatment).

### Polyclonal antibody development

CD200 and 41BB FAST proteins were digested overnight with TEV protease (Sigma # T4455) at 4 °C in PBS. The cleaved linker and 6x-His tag were then removed using a His SpinTrap kit (GE Healthcare # 28-9321-71). Digested proteins in PBS were diluted 1:1 in Squalvax (Oz Biosciences # SQ0010) to a final concentration of 0.1 μg/μL and was mixed using interlocked syringes to form an emulsion. Immunization of BALB/c mice for antibody production was approved by the University of Tasmania Animal Ethics Committee (# A0014680). Pre-immune sera were collected prior to subcutaneous immunization with at least 50 μL of the emulsion. On day 14 post-immunization the mice were boosted using a similar procedure. On day 50 the mice received a booster with proteins in IFAVax (Oz Biosciences # IFA0050); mice immunized with CD200 again received subcutaneous injections, whereas 41BB mice received subcutaneous and intraperitoneal injections. Pre-immune and sera collected after 3X immunizations were then tested by flow cytometry against CHO cells expressing either 41BB or CD200. CHO cells were prepared as described above and 2×10^5^ cells were incubated with mouse serum diluted 1:200 in PBS for 30 minutes at 4 °C. The cells were then washed 2X and stained with 50 μL of anti-mouse IgG AlexaFluor 647 diluted 1:1000 in FACS buffer. The cells were then washed 2X, stained with DAPI to identify live cells, and analyzed on a Cyan ADP flow cytometer (Figure 5C). CD200 and 41BB expression on DFT cells was tested using a procedure similar to the CHO cell staining, except the sera used was collected after 4X immunizations and was diluted 1:500 and analyzed on the BD FACSCanto II (Figure 5D).

### Purification of antibodies from normal mouse serum (NMS) and anti-CD200 serum

Approximately 200 μL of normal mouse serum or anti-CD200 serum day 157 (after 4X immunizations) were purified using a protein G SpinTrap (GE Healthcare # 28-4083-47) according to the manufacturer’s instrutions. Serum was diluted 1:1 with 20 mM sodium phosphate, pH 7.0 binding buffer, eluted with 0.1 M glycine-HCl, pH 2.7, and the pH was neutralized with 0.1 M glycine-HCl, pH 2.7. The eluted antibodies were then concentrated using an Amicon Ultra 0.5 centrifugal until (Merck # UFC500308) by centrifuging at 14,000 RCF for 30 minutes at 4 °C and then washing the antibodies with 400 μL of PBS twice. The protein concentration was then quantified on a Nanodrop spectrophotometer at 280 nm using the extinction coefficients for IgG.

### Testing CD200 expression on DFT cells that overexpress CD200R1

50,000 DFT cells/well were aliquoted into u-bottom 96-well plates and washed with 200 μL of cold FACS buffer. Purified polyclonal anti-CD200 was diluted to 2.5 μg/mL in cold FACS buffer and the cells in appropriate wells were resuspend in 100 μL/well (0.25 μg/well) diluted antibody; wells that did not receive antibody were resuspended in 100 μL of FACS buffer. The cells were incubated on ice for 20 minutes and then washed with 200 μL of FACS buffer. Whilst incubating, anti-mouse IgG-AF647 was diluted to 1 μg/mL in cold FACS buffer and then used to resuspend cells in the appropriate wells. The plates were incubated on ice for 20 minutes, then washed with 100 μL of cold FACS buffer. The cells were then resuspended in 200 μL of FACS fix and incubated on a rocking platform at room temp for 15 minutes. The cells were then centrifuged 500 RCF for 3 minutes at 4 °C, resuspended in 200 μL FACS buffer and stored at 4°C until they were analyzed on a FACSCanto II (n=2/treatment).

### Isolation of devil peripheral blood mononuclear cells

Blood collection from Tasmanian devils was approved by the University of Tasmania Animal Ethics Committee (permit # A0014599) and the Tasmanian Department of Primary Industries, Parks, Water and Environment (DPIPWE). Blood was collected from the jugular vein and stored in EDTA tubes for transport to the lab. Blood was processed within three hours by diluting 1:1 with serum-free RPMI and then layering onto Histopaque (Sigma # 10771) before centrifuging at 400 RCF for 30 minutes. The interface containing the peripheral blood mononuclear cells was then collected using a transfer pipette, diluted with 50 mL of serum-free RPMI and centrifuged for 5 minutes at 500 RCF. Cells were washed with again with cRF10 and then either used fresh or stored at −80 °C until further use.

### Detecting DFT2 cells in PBMC using CD200

Frozen devil PBMC were thawed and cultured in cRF10 at 35 °C with 5% CO_2_ for 2 hours, cells were then washed in FACS buffer, counted and 3×10^5^ PBMC cells used per sample. DFT2.JV cells were removed from culture flasks, counted, and 2×10^5^ cells used per sample. Samples were incubated with 50 μL normal goat serum (Thermo Cat # 01-6201) diluted 1:200 in FACS buffer for 15 minutes at 4 °C, 50 μL of anti-CD200 serum diluted 1:100 was added (1:200 final) for 30 minutes at 4 °C. Cells were then washed 2X and stained with 50 μL of anti-mouse IgG AlexaFluor 647 diluted to 1 μg/mL in FACS buffer for 30 minutes at 4 °C. The cells were then washed 2X, stained with DAPI (Sigma Cat # D9542) to identify live cells, and analyzed on the BD FACSCanto II. PBMC and DFT cells were run separately then PBMC and DFT2 mixed at a ratio of 10:1 by volume for the combined samples (n=1/treatment) (Figure S4A). The experiment was repeated (n=1/treatment), except that PBMCs and DFT cells were mixed at a 5:1 ratio (Figure S4B).

### Staining of DFT cells in devil whole blood

DFT1.C5065 and DFT2.JV cells were labelled with 5 μM CellTrace violet (CTV) and cultured for three days at 37 °C. On the day of the assays peripheral blood from one devil was collected and stored at ambient temp for less than three hours. 100 μL of whole blood was aliquoted into 15 mL tubes and stored at ambient temperature whilst DFT cells were prepared. The media on CTV-labeled DFT cells were decanted and the cells were detached from the flask by incubating in 2.5 mL of TrypLE Select for 5 minutes at 37 °C. The cells were washed with cRF10, resuspended in cRF10, and counted. DFT cells were then diluted to 1×10^4^ cells/mL in cRF10 and 100 μL were aliquoted into appropriate 15 mL tubes containing 100 μL of whole blood. 1 μL of purified anti-CD200 (0.5 μg/tube) was diluted into the appropriate tubes and incubated for 15 minutes at ambient temperature. Next, 0.5 μg/tube of anti-mouse IgG AF647 was added to each tube. Note: 0.5 μL (0.5 μg) of concentrated secondary antibody was accidentally added directly to the tube for the data shown in the top row and middle column of Figure S5A; for all other tubes the secondary antibody was diluted 1:20 in PBS and 10 μL was added to each tube. The cells were then incubated for 15 minutes at ambient temperature. The cells were then diluted in 1 mL ammonium chloride red blood cell (RBC) lysis buffer (150 mM NH_4_Cl, 10 mM KHCO_3_, 0.1 mM EDTA disodium (Na_2_-2H_2_O)) and mixed immediately gently pipetting five times. The cells were incubated at ambient temperature for 10 minutes and then diluted with 5 mL of PBS and centrifuged 500 RCF for 3 minutes. Some tubes contained residual RBCs, so the pellet was vigorously resuspended in 5 mL of RBC lysis buffer, incubated for 5 minutes, diluted with 5 mL of cold FACS buffer, and centrifuged 500 RCF for 3 minutes. The cells were then resuspended in 250 μL of FACS buffer and stored on ice until analysis on a Beckman Coulter MoFlo Astrios (n=1/treatment). Data were analyzed in FCS Express version 6 (Figure S5).

The experiment above was repeated with the following modifications. DFT cells were labelled with 5 μM of CFSE and incubated for two days at 37 °C. On the day of the assays fresh blood was collected from two devils. Purified anti-CD200 and NMS were labeled with Zenon mouse IgG AF647 (ThermoFisher # Z25008) and blocked with the Zenon blocking agent. 1×10^4^ CFSE-labeled DFT cells were diluted directly into 100 μL of whole blood in 15 mL tubes and 12 μL (2 μL antibody, 5 μL labeling agent, 5 μL blocking agent) of Zenon AF647-labeled purified NMS or anti-CD200 were added directly to the cells. The cells were incubated for 30 minutes at ambient temperature. The cells were then gently resuspended in 2.5 mL of RBC lysis buffer and incubated for 10 minutes at ambient temperature. The cells were diluted with 10 mL of PBS and centrifuged 500 RCF for 3 minutes. The cells were resuspended in 1.5 mL of RBC lysis buffer and incubated for another 10 minutes to lyse residual RBCs. The tubes were then resuspended in 9 mL of cRF10 and centrifuged 500 RCF for 3 minutes. The cells were resuspended in 350 μL of cold FACS buffer containing 200 ng/mL of DAPI and stored on ice until analysis on a Beckman Coulter MoFlo Astrios (n=1/treatment for n = 2 devils) (Figure 6).

TABLES (See supplementary Materials)

## ACKNOWLEDGMENTS

We wish to thank Ginny Ralph for her ongoing care of Tasmanian devils, and the Bonorong Wildlife Sanctuary for providing access to Tasmanian devils, and Ruth Pye for providing care for devils and collecting blood samples. We would like to thank John Hayball and Georgina Kalodimos for advice and assistance, Mahalia Kingsley and Nirdesh Poudel for plasmid construction, Nick Blackburn for bioinformatics assistance, Lynn Corcoran for the devil IgG plasmid, James Murphy for a devil IL-15 plasmid, Emily Flies for editing the manuscript, and Kay Holekamp for her comments on the manuscript. Funding: ARC DECRA grant # DE180100484, ARC Linkage grant # LP0989727, ARC Discovery grant # DP130100715, Morris Animal Foundation Grant-in-Aid # D14ZO-410, University of Tasmania Foundation Dr Eric Guiler Tasmanian Devil Research Grant through funds raised by the Save the Tasmanian Devil Appeal (2013, 2015, 2017, 2018), and Entrepreneurs’ Programme - Research Connections grant with Nexvet Australia Pty. Ltd. # RC50680.

## AUTHOR CONTRIBUTIONS

ASF designed the study; ALP, ASF, CEBO, PRL, and PRM developed the technology; ASF, CEBO, PRL, PRM, JMD, and TLP performed the experiments; ALP performed bioinformatic analyses; ALP, ASF, JMD and PRL created the figures; ALP, ASF, PRL, JMD, TLP, and GMW analyzed the data; ASF wrote the manuscript and all authors edited the manuscript.

## AVAILABILITY OF DATA AND MATERIALS

All data are included with the paper. The FAST base vectors (pAF92.3 pAF112.7, pAF123.1, pAF138.7, pAF139.2, pAF160.1, pAF161.3, pAF163.1, pAF164.3, pAF197) have been submitted to Addgene for distribution c(deposit # 77504).

## SUPPLEMENTARY INFORMATION

### SUPPLEMENTARY MATERIALS AND METHODS

**Figure S1.**
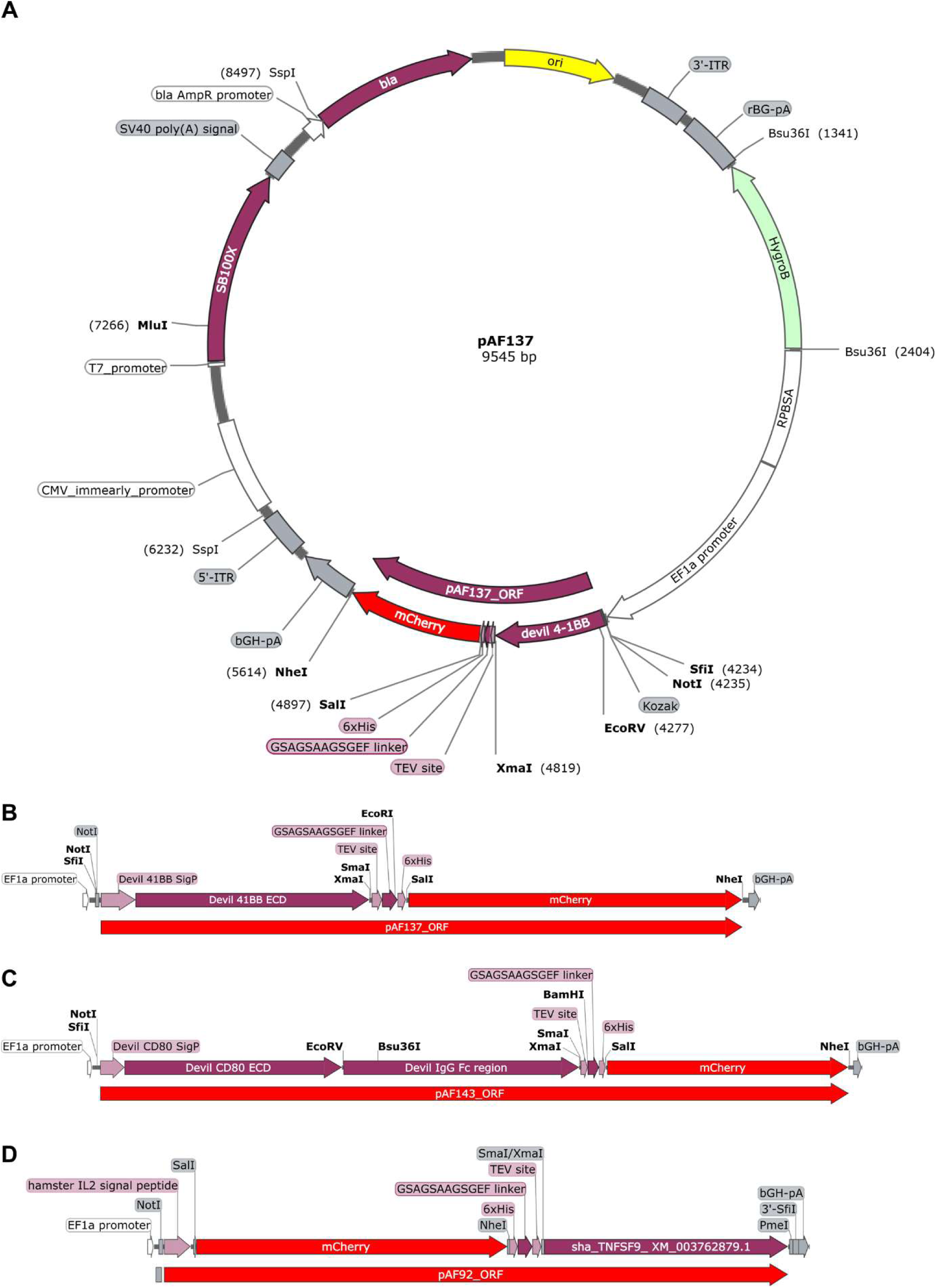
FAST protein base vectors. (A) CMV promotor and SB100X transposase were inserted into a Sleeping Beauty vector (shown here with 41BB-mCherry FAST protein cassette pAF137). (B) Type I FAST protein cassette. The native signal peptide (SigP) and predicted extracellular domain (ECD) were used for all type I FAST proteins. The devil 41BB SigP-ECD was fused to mCherry via a TEV cleavage site (ENLYFQG), linker peptide (GSAGSAAGSGEF), and a 6x-His tag (HHHHHH). Restriction digest sites were included at the 5’ and 3’ ends of the gene-of-interest and the fluorescent protein to facilitate swapping of genes and fluorescent proteins. The human EF1α promotor is upstream and bovine growth hormone (bGH) poly(A) tail is downstream of the open reading frame. (C) Type I FAST protein cassette with Fc tag. This vector was the same as the type I FAST vectors, except that the Fc fragment of devil IgG was inserted between the gene-of-interest (e.g. CD80) and the TEV cleavage site. (D) Type II FAST protein cassette. To increase the probability of efficient secretion of type II FAST proteins from Chinese hamster ovary (CHO) cells, we used the hamster IL-2 signal peptide at the N-terminus of the protein, including a SalI restriction site, followed by mCherry, an NheI restriction site, 6x-His-tag, linker peptide, TEV cleavage site, XmaI/SmaI restriction site, the gene-of-interest, and a PmeI restriction site following the stop codon.

**Figure S2.**
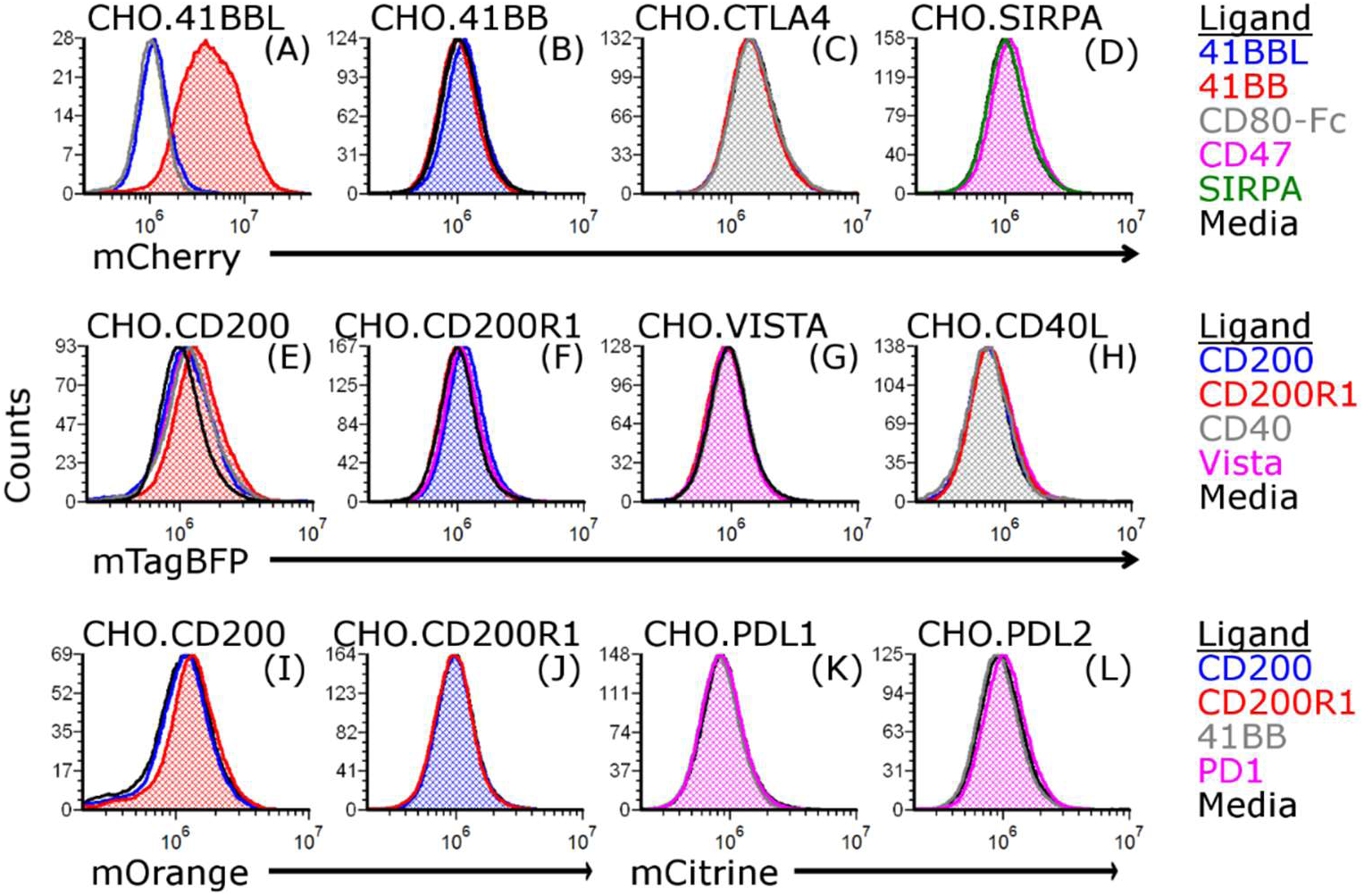
Screening FAST protein supernatants. Staining was performed using supernatants from FAST protein CHO cell lines; supernatants were collected 11 days post-transfection. Supernatants from FAST cell lines were incubated with CHO cells expressing full-length devil proteins at room temperature for 30 minutes before washing and fixing for flow cytometric analysis. Chloroquine was not used during these incubation steps.

**Figure S3.**
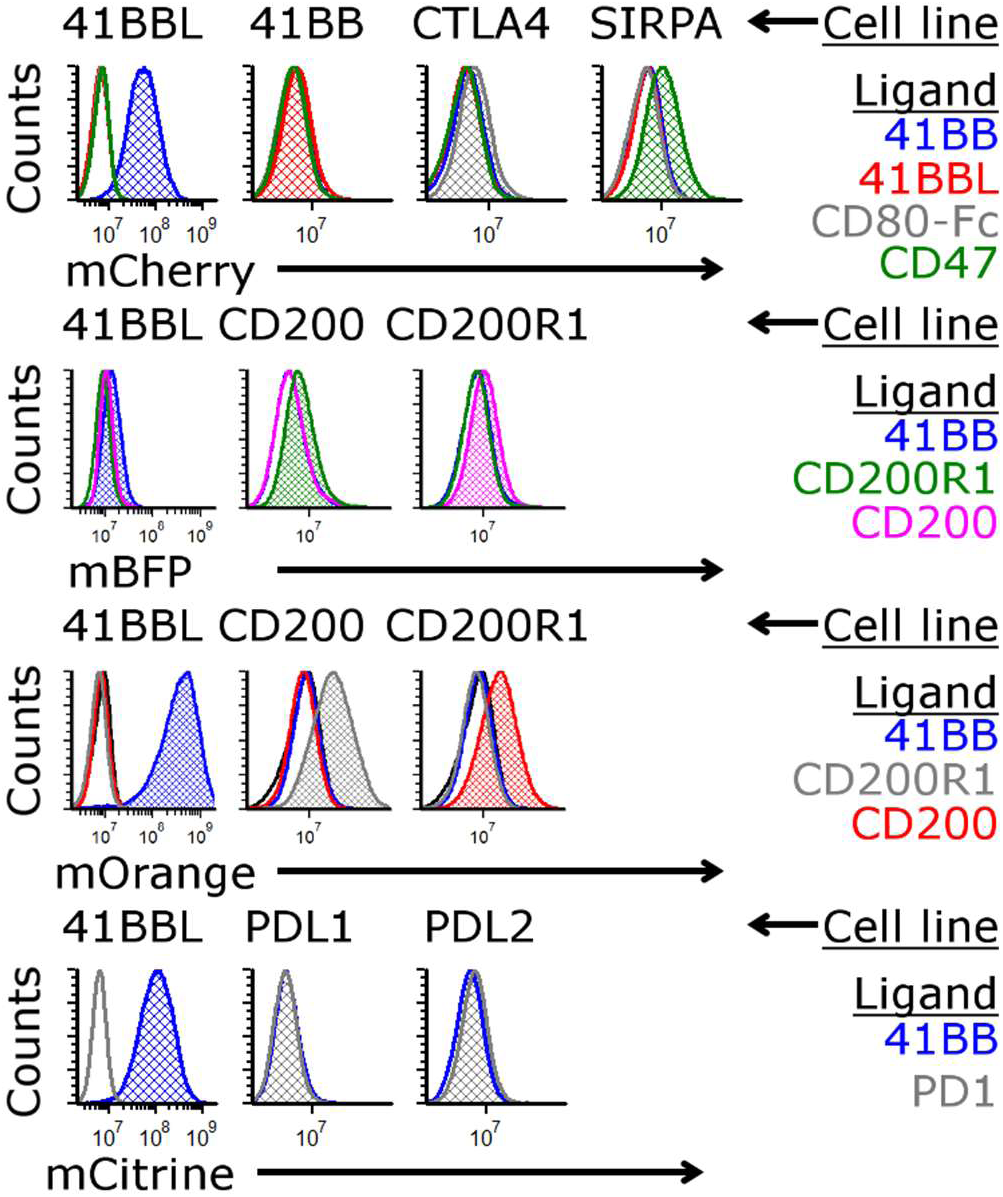
Screening FAST protein supernatants. Staining was performed using supernatants from FAST protein CHO cell lines; supernatants were collected around 2-3 weeks post-transfection and stored for 2 months at 4 °C. Supernatants from FAST cell lines were diluted 1:1 with 100 μM chloroquine (50 μM final concentration) in cRF5 without phenol red incubated with CHO cells expressing full-length devil proteins at room temperature for 60 minutes before washing and fixing for flow cytometric analysis.

**Figure S4.**
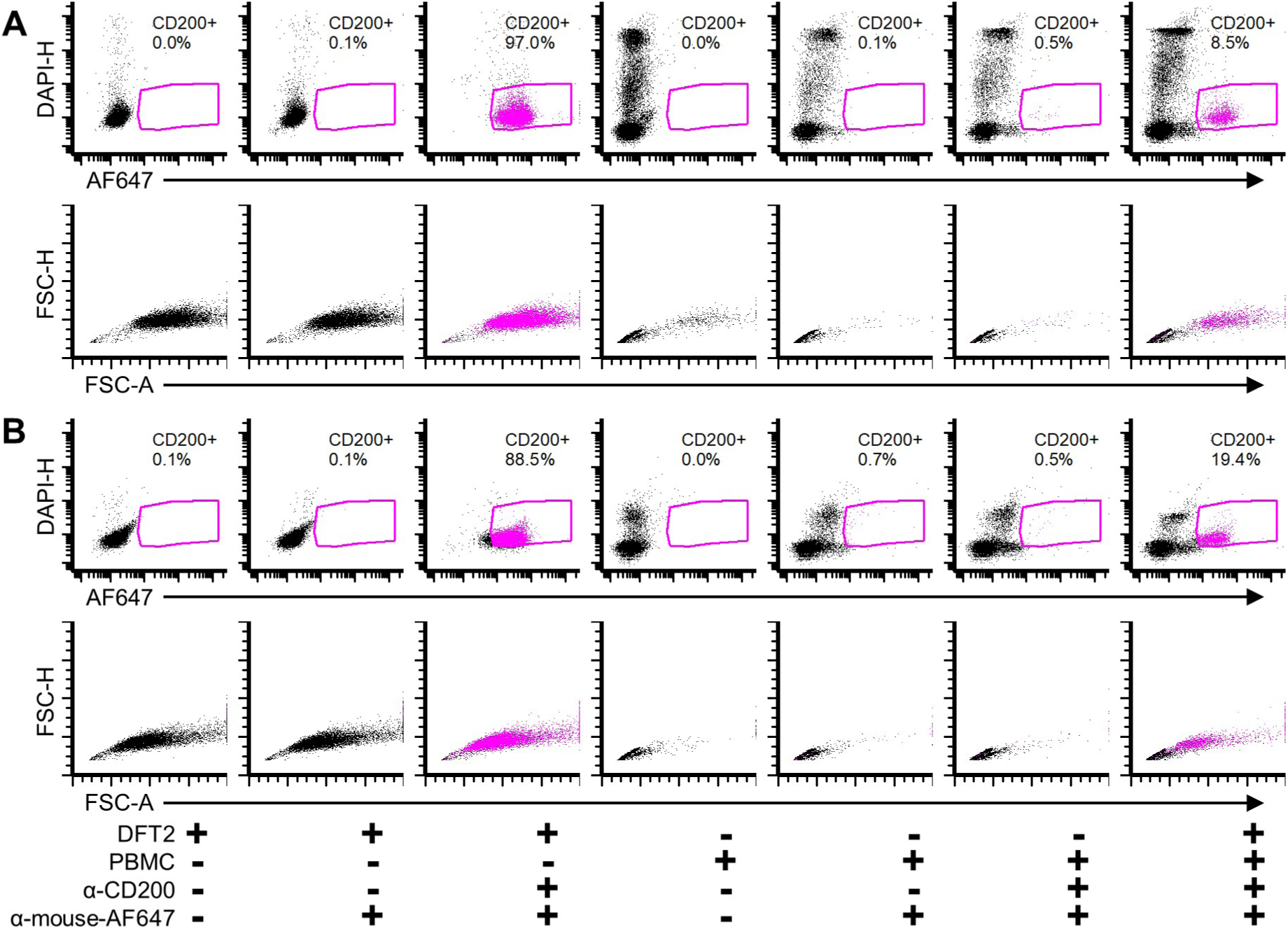
CD200 identifies DFT2 cells in PBMCs. (A) PBMCs isolated on histopaque, frozen, thawed, and cultured for 2 hours at 37 °C prior to use. DFT2 cells and PBMCs were stained separately by blocking with normal goat serum and then staining with anti-CD200 serum and anti-mouse IgG Alexa-647. Cells were then washed, stained with DAPI, and analyzed on a BD FACSCanto II. The stained DFT2 cells and PBMCs were then mixed at 1:10 to see if gating and CD200 signal could be used to distinguish DFT2 cells from PBMCs (n=1/treatment). (B) The procedure used for A was repeated except that DFT2 cells and PBMCs were mixed at 1:5.

**Figure S5.**
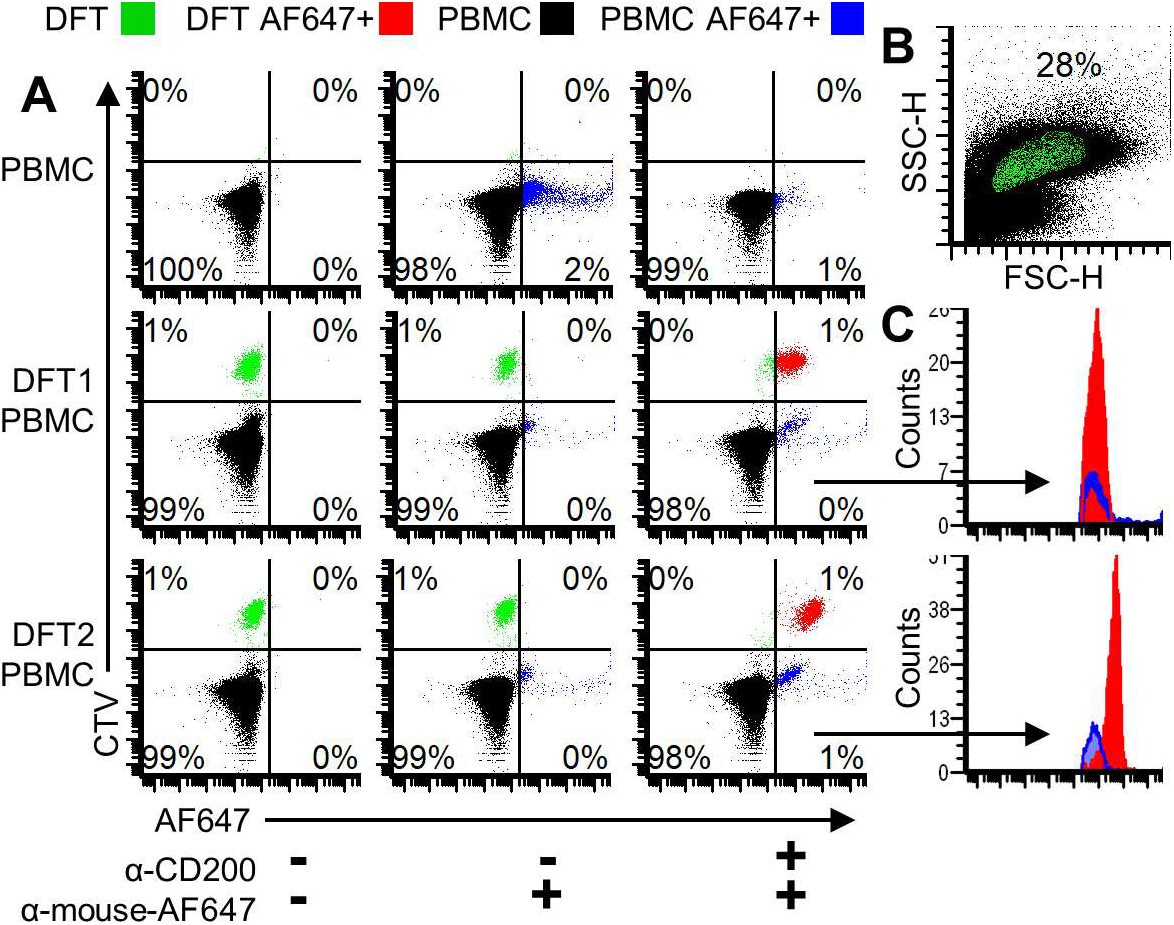
CD200 identifies DFT cells in whole blood. Color dot plots showing DFT cells in green (CellTrace Violet®), PBMCs in black, DFT Alexa Fluor 647+ (AF647) cells in red, and PBMC AF647+ in blue. (A) The top row shows unmixed PBMCs. The middle row and bottom row show DFT1.C5065 and DFT2.JV cells, respectively, mixed with PBMCs. Alexa Fluor 647+ DFT (red) and PBMC (blue) are in the right quadrants. (B) Forward-and side-scatter plot of DFT.JV cells mixed with PBMCs. Backgating of CFSE+ cells was used to create a forward-scatter by side-scatter gate thaat was used as the parent gate for all data shown here. (C) Histogram overlays to highlight AF647+ (right quadrants) from DFT1-PBMC and DFT2-PBMC mixtures. Cells were analyzed on the Beckman-Coulter MoFlo Astrios.

**Table S1. Search terms for mammalian immune research studies.**

**Table S2. Summary and genes and plasmids.**

**Table S3. Vector backbones and insert cassette components.**

**Table S4. Primer sequences and gBlocks for plasmid assembly.**

**Table S5. gBlocks used for assembling plasmids.**

## REFERENCES

1. Albuquerque TAF, Drummond do Val L, Doherty A, de Magalhães JP. From humans to hydra: patterns of cancer across the tree of life. Biol Rev 2018; 93: 1715–1734.

2. Abu-Helil B, Van Der Weyden L. Metastasis in the wild: investigating metastasis in non-laboratory animals. Clin Exp Metastasis 2019. doi:10.1007/s10585-019-09956-3.

3. Olds JE, Burrough ER, Fales-Williams AJ, Lehmkuhl A, Madson D, Patterson AJ et al. Retrospective Evaluation of Cases of Neoplasia in a Captive Population of Egyptian Fruit Bats (Rousettus Aegyptiacus). J Zoo Wildl Med 2015; 46: 325–332.

4. Chu PY, Zhuo YX, Wang FI, Jeng CR, Pang VF, Chang PH et al. Spontaneous neoplasms in zoo mammals, birds, and reptiles in Taiwan - A 10-year survey. Anim Biol 2012; 62: 95–110.

5. Deus A De, Alves F, Siqueira DB De, Rameh-de-albuquerque LC. Breast Carcinoma with Pulmonary Metastasis in Armadillo (Eupharactus sexcinctus). Acta Sci Vet 2018; 46: 1–5.

6. Kruse TN, Garner MM, Bonar CJ. A Retrospective Study of Pathologic Findings in Captive Rock Hyrax (Procavia Capensis) in the United States. J Zoo Wildl Med 2015; 46: 798–805.

7. Effron M, Griner L, Benirschke K. Nature and rate of neoplasia found in captive wild mammals, birds, and reptiles at necropsy. J Natl Cancer Inst 1977; 59: 185–198.

8. Ratcliffe HL. Incidence and nature of tumors in captive wild mammals and birds. Am J Cancer 1933; 17: 116–135.

9. Priester WA, Mantel N. Occurrence of tumors in domestic animals. Data from 12 United States and Canadian colleges of veterinary medicine. J Natl Cancer Inst 1971; 47: 1333–1344.

10. Howlader N, Noone A, Krapcho M, Miller D, Brest A, Yu M et al. SEER Cancer Statistics Review, 1975-2016. Bethesda, MD, United States, 2016.

11. Griner LA. Neoplasms in Tasmanian devils (Sarcophilus harrisii). J Natl Cancer Inst 1979; 62: 589–595.

12. Peck SJ, Michael SA, Knowles G, Davis A, Pemberton D. Causes of mortality and severe morbidity requiring euthanasia in captive Tasmanian devils (Sarcophilus harrisii) in Tasmania. Aust Vet J 2019; 97: 89–92.

13. Pearse A-M, Swift K. Allograft theory: transmission of devil facial-tumour disease. Nature 2006; 439: 549.

14. Pye RJ, Pemberton D, Tovar C, Tubio JMC, Dun KA, Fox S et al. A second transmissible cancer in Tasmanian devils. Proc Natl Acad Sci 2016; 113: 374–379.

15. Murgia C, Pritchard JK, Kim SY, Fassati A, Weiss RA. Clonal Origin and Evolution of a Transmissible Cancer. Cell 2006; 126: 477–487.

16. Fleming JM, Creevy KE, Promislow DEL. Mortality in North American Dogs from 1984 to 2004: An Investigation into Age-, Size-, and Breed-Related Causes of Death. J Vet Intern Med 2011; 25: 187–198.

17. Lazenby BT, Tobler MW, Brown WE, Hawkins CE, Hocking GJ, Hume F et al. Density trends and demographic signals uncover the long-term impact of transmissible cancer in Tasmanian devils. J Appl Ecol 2018; 55: 1368–1379.

18. James S, Jennings G, Kwon YM, Stammnitz M, Fraik A, Storfer A et al. Tracing the rise of malignant cell lines: distribution, epidemiology and evolutionary interactions of two transmissible cancers in Tasmanian devils. Evol Appl 2019;: eva.12831.

19. Siddle H V., Kreiss A, Tovar C, Yuen CK, Cheng Y, Belov K et al. Reversible epigenetic down-regulation of MHC molecules by devil facial tumour disease illustrates immune escape by a contagious cancer. Proc Natl Acad Sci 2013; 110: 5103–8.

20. Yoshihama S, Roszik J, Downs I, Meissner TB, Vijayan S, Chapuy B et al. NLRC5/MHC class I transactivator is a target for immune evasion in cancer. Proc Natl Acad Sci 2016; 113: 5999–6004.

21. Caldwell A, Coleby R, Tovar C, Stammnitz MR, Mi Kwon Y, Owen RS et al. The newly-arisen devil facial tumour disease 2 (DFT2) reveals a mechanism for the emergence of a contagious cancer. Elife 2018; 7. doi:10.7554/eLife.35314.

22. Stammnitz MR, Coorens THH, Gori KC, Hayes D, Fu B, Wang J et al. The Origins and Vulnerabilities of Two Transmissible Cancers in Tasmanian Devils. Cancer Cell 2018; 33: 607–619.e15.

23. Topalian SL, Hodi FS, Brahmer JR, Gettinger SN, Smith DC, McDermott DF, et al. Safety, Activity, and Immune Correlates of Anti–PD-1 Antibody in Cancer. N Engl J Med 2012; 366: 2443–2454.

24. Larkin J, Chiarion-Sileni V, Gonzalez R, Grob JJ, Cowey CL, Lao CD et al. Combined Nivolumab and Ipilimumab or Monotherapy in Untreated Melanoma. N Engl J Med 2015; 373: 23–34.

25. Flies AS, Bruce Lyons A, Corcoran LM, Papenfuss AT, Murphy JM, Knowles GW et al. PD-L1 is not constitutively expressed on tasmanian devil facial tumor cells but is strongly upregulated in response to IFN-γ and can be expressed in the tumor microenvironment. Front Immunol 2016; 7: 581.

26. Ong CEB, Lyons AB, Woods GM, Flies AS. Inducible IFN-γ Expression for MHC-I Upregulation in Devil Facial Tumor Cells. Front Immunol 2018; 9: 3117.

27. Flies AS, Blackburn NB, Lyons AB, Hayball JD, Woods GM. Comparative analysis of immune checkpoint molecules and their potential role in the transmissible tasmanian devil facial tumor disease. Front Immunol 2017; 8: 513.

28. World Health Organization (WHO). WHO R&D Blueprint for action to prevent epidemics. World Health Organization, 2016 http://www.who.int/blueprint/en/ (accessed 1 May2018).

29. Shaner NC, Campbell RE, Steinbach PA, Giepmans BNG, Palmer AE, Tsien RY. Improved monomeric red, orange and yellow fluorescent proteins derived from Discosoma sp. red fluorescent protein. Nat Biotechnol 2004; 22: 1567–72.

30. Cabantous S, Terwilliger TC, Waldo GS. Protein tagging and detection with engineered self-assembling fragments of green fluorescent protein. Nat Biotechnol 2005; 23: 102– 107.

31. Cha HJ, Dalal NN, Bentley WE. Secretion of human interleukin-2 fused with green fluorescent protein in recombinant Pichia pastoris. Appl Biochem Biotechnol 2005; 126: 1–11.

32. Duellman T, Burnett J, Yang J. Quantitation of secreted proteins using mCherry fusion constructs and a fluorescent microplate reader. Anal Biochem 2015; 473: 34–40.

33. Kammertoens T, Friese C, Arina A, Idel C, Briesemeister D, Rothe M et al. Tumour ischaemia by interferon-γ resembles physiological blood vessel regression. Nature 2017; 545: 98–102.

34. Patchett AL, Wilson R, Charlesworth JC, Corcoran LM, Papenfuss AT, Lyons AB et al. Transcriptome and proteome profiling reveals stress-induced expression signatures of imiquimod-treated Tasmanian devil facial tumor disease (DFTD) cells. Oncotarget 2018; 9: 15895–15914.

35. Patchett AL, Coorens THH, Darby J, Wilson R, McKay MJ, Kamath KS et al. Two of a kind: transmissible Schwann cell cancers in the endangered Tasmanian devil (Sarcophilus harrisii). Cell Mol Life Sci 2019;: 1–12.

36. Seglen PO, Grinde B, Solheim AE. Inhibition of the Lysosomal Pathway of Protein Degradation in Isolated Rat Hepatocytes by Ammonia, Methylamine, Chloroquine and Leupeptin. Eur J Biochem 1979; 95: 215–225.

37. Qureshi OS, Zheng Y, Nakamura K, Attridge K, Manzotti C, Schmidt EM et al. Trans-endocytosis of CD80 and CD86: a molecular basis for the cell extrinsic function of CTLA-4. Science 2011; 332: 600–603.

38. van den Bremer ET, Beurskens FJ, Voorhorst M, Engelberts PJ, de Jong RN, van der Boom BG et al. Human IgG is produced in a pro-form that requires clipping of C-terminal lysines for maximal complement activation. MAbs 2015; 7: 672–680.

39. Patchett AL, Latham R, Brettingham-Moore KH, Tovar C, Lyons AB, Woods GM. Toll-like receptor signaling is functional in immune cells of the endangered Tasmanian devil. Dev Comp Immunol 2015; 53: 123–133.

40. Love JE, Thompson K, Kilgore MR, Westerhoff M, Murphy CE, Papanicolau-Sengos A et al. CD200 Expression in Neuroendocrine Neoplasms. Am J Clin Pathol 2017; 148: 236–242.

41. Murchison EP, Tovar C, Hsu A, Bender HS, Kheradpour P, Rebbeck CA et al. The Tasmanian devil transcriptome reveals schwann cell origins of a clonally transmissible cancer. Science 2010; 327: 84–87.

42. Haile ST, Bosch JJ, Agu NI, Zeender AM, Somasundaram P, Srivastava MK et al. Tumor cell programmed death ligand 1-mediated T cell suppression is overcome by coexpression of CD80. J Immunol 2011; 186: 6822–9.

43. Sugiura D, Maruhashi T, Okazaki I, Shimizu K, Maeda TK, Takemoto T et al. Restriction of PD-1 function by cis-PD-L1/CD80 interactions is required for optimal T cell responses. Science 2019;: eaav7062.

44. Moertel CL, Xia J, LaRue R, Waldron NN, Andersen BM, Prins RM et al. CD200 in CNS tumor-induced immunosuppression: The role for CD200 pathway blockade in targeted immunotherapy. J Immunother Cancer 2014; 2: 46.

45. Challagundla P, Medeiros LJ, Kanagal-Shamanna R, Miranda RN, Jorgensen JL. Differential Expression of CD200 in B-Cell Neoplasms by Flow Cytometry Can Assist in Diagnosis, Subclassification, and Bone Marrow Staging. AJCP / Orig Artic Am J Clin Pathol 2014; 142: 837–844.

46. Spacek M, Karban J, Radek M, Babunkova E, Kvasnicka J, Jaksa R et al. CD200 Expression Improves Differential Diagnosis Between Chronic Lymphocytic Leukemia and Mantle Cell Lymphoma. Blood 2014; 124.

47. Saksena A, Yin CC, Xu J, Li J, Zhou J, Wang SA et al. CD23 expression in mantle cell lymphoma is associated with CD200 expression, leukemic non-nodal form, and a better prognosis. Hum Pathol 2019; 89: 71–80.

48. Loh R, Hayes D, Mahjoor A, O’Hara A, Pyecroft S, Raidal S. The immunohistochemical characterization of devil facial tumor disease (DFTD) in the Tasmanian Devil (Sarcophilus harrisii). Vet Pathol 2006; 43: 896–903.

49. Yao S, Zhu Y, Zhu G, Augustine M, Zheng L, Goode DJ et al. B7-h2 is a costimulatory ligand for CD28 in human. Immunity 2011; 34: 729–740.

50. Wang J, Sanmamed MF, Datar I, Su TT, Ji L, Sun J et al. Fibrinogen-like Protein 1 Is a Major Immune Inhibitory Ligand of LAG-3. Cell 2018; 0. doi:10.1016/J.CELL.2018.11.010.

51. Aricescu AR, Lu W, Jones EY. A time-and cost-efficient system for high-level protein production in mammalian cells. Acta Crystallogr Sect D Biol Crystallogr 2006; 62: 1243– 1250.

52. Consortium SG, Biologiques A et F des M, Center BSG, Consortium CSG, Innovation IC for S and F, Center ISP et al. Protein production and purification. Nat Methods 2008; 5: 135.

53. Jayapal KP, Wlaschin KF, Hu WS, Yap MGS. Recombinant protein therapeutics from CHO cells - 20 years and counting. Chem Eng Prog 2007; 103: 40–47.

54. Atfy M. CD200 Suppresses the Natural Killer Cells and Decreased its Activity in Acute Myeloid Leukemia Patients. J Leuk 2015; 3.https://www.omicsonline.org/open-access/cd200-suppresses-the-natural-killer-cells-and-decreased-its-activity-in-acutemyeloid-leukemia-patients-2329-6917-1000190.pdf (accessed 18 Apr2019).

55. Coles SJ, Man S, Hills R, Wang EC, Burnett A, Darley RL et al. Over-Expression of CD200 In Acute Myeloid Leukemia Mediates the Expansion of Regulatory T-Lymphocytes and Directly Inhibits Natural Killer Cell Tumor Immunity. Blood 2015; 116: 491.

56. Coles SJ, Wang ECY, Man S, Hills RK, Burnett AK, Tonks A et al. CD200 expression suppresses natural killer cell function and directly inhibits patient anti-tumor response in acute myeloid leukemia. Leukemia 2011; 25: 792–9.

57. Gorczynski RM, Chen Z, Khatri I, Yu K. Graft-infiltrating cells expressing a CD200 transgene prolong allogeneic skin graft survival in association with local increases in Foxp3 +Treg and mast cells. Transpl Immunol 2011; 25: 187–193.

58. Gorczynski RM, Chen Z, He W, Khatri I, Sun Y, Yu K et al. Expression of a CD200 transgene is necessary for induction but not maintenance of tolerance to cardiac and skin allografts. J Immunol 2009; 183: 1560–1568.

59. Gorczynski L, Chen Z, Hu J, Kai Y, Lei J, Ramakrishna V et al. Evidence That an OX-2-Positive Cell Can Inhibit the Stimulation of Type 1 Cytokine Production by Bone Marrow-Derived B7-1 (and B7-2)-Positive Dendritic Cells. J Immunol 1999; 162: 774– 781.

60. Harding J, Vintersten-Nagy K, Shutova M, Yang H, Tang JK, Massumi M et al. Induction of long-term allogeneic cell acceptance and formation of immune privileged tissue in immunocompetent hosts. Cold Spring Harbor Laboratory, 2019 doi:10.1101/716571.

61. Jenmalm MC, Cherwinski H, Bowman EP, Phillips JH, Sedgwick JD. Regulation of Myeloid Cell Function through the CD200 Receptor. J Immunol 2014; 176: 191–199.

62. Tovar C, Pye RJ, Kreiss A, Cheng Y, Brown GK, Darby J et al. Regression of devil facial tumour disease following immunotherapy in immunised Tasmanian devils. Sci Rep 2017; 7: 43827.

63. Pye R, Patchett A, McLennan E, Thomson R, Carver S, Fox S et al. Immunization strategies producing a humoral IgG immune response against devil facial tumor disease in the majority of Tasmanian devils destined for wild release. Front Immunol 2018; 9: 259.

64. Olin MR, Ampudia-Mesias E, Pennell CA, Sarver A, Chen CC, Moertel CL et al. Treatment combining CD200 immune checkpoint inhibitor and tumor-lysate vaccination after surgery for pet dogs with high-grade glioma. Cancers (Basel) 2019; 11: 137.

65. Zhang S, Phillips JH. Identification of tyrosine residues crucial for CD200R-mediated inhibition of mast cell activation. J Leukoc Biol 2005; 79: 363–368.

66. Hatherley D, Lea SM, Johnson S, Barclay AN. Structures of CD200/CD200 receptor family and implications for topology, regulation, and evolution. Structure 2013; 21: 820– 832.

67. Luo ZX, Yuan CX, Meng QJ, Ji Q. A Jurassic eutherian mammal and divergence of marsupials and placentals. Nature 2011; 476: 442–445.

68. Wong KK, Khatri I, Shaha S, Spaner DE, Gorczynski RM. The role of CD200 in immunity to B cell lymphoma. J Leukoc Biol 2010; 88: 361–372.

69. Caserta S, Nausch N, Sawtell A, Drummond R, Barr T, MacDonald AS et al. Chronic infection drives expression of the inhibitory receptor CD200R, and its ligand CD200, by mouse and human CD4 T cells. PLoS One 2012; 7: e35466.

70. Foster-Cuevas M, Westerholt T, Ahmed M, Brown MH, Barclay AN, Voigt S. Cytomegalovirus e127 protein interacts with the inhibitory CD200 receptor. J Virol 2011; 85: 6055–9.

71. Foster-Cuevas M, Wright GJ, Puklavec MJ, Brown MH, Barclay AN. Human herpesvirus 8 K14 protein mimics CD200 in down-regulating macrophage activation through CD200 receptor. J Virol 2004; 78: 7667–76.

72. Alves JM, Carneiro M, Cheng JY, Matos AL de, Rahman MM, Loog L, et al. Parallel adaptation of rabbit populations to myxoma virus. Science 2019;: eaau7285.

73. Barclay AN, Hatherley D. The Counterbalance Theory for Evolution and Function of Paired Receptors. Immunity 2008; 29: 675–678.

74. Tovar C, Obendorf D, Murchison EP, Papenfuss AT, Kreiss A, Woods GM. Tumor-specific diagnostic marker for transmissible facial tumors of tasmanian devils: Immunohistochemistry studies. Vet Pathol 2011; 48: 1195–1203.

75. Marx V. Calling the next generation of affinity reagents. Nat Methods 2013; 10: 829–833.

76. Grabherr MG, Haas BJ, Yassour M, Levin JZ, Thompson DA, Amit I et al. Full-length transcriptome assembly from RNA-Seq data without a reference genome. Nat Biotechnol 2011; 29: 644–52.

77. Kowarz E, Löscher D, Marschalek R. Optimized Sleeping Beauty transposons rapidly generate stable transgenic cell lines. Biotechnol J 2015; 10: 647–653.

78. Mátés L, Chuah MKL, Belay E, Jerchow B, Manoj N, Acosta-Sanchez A et al. Molecular evolution of a novel hyperactive Sleeping Beauty transposase enables robust stable gene transfer in vertebrates. Nat Genet 2009; 41: 753–761.

79. Waldo GS, Standish BM, Berendzen J, Terwilliger TC. Rapid protein-folding assay using green fluorescent protein. Nat Biotechnol 1999; 17: 691–5.

80. Kozak M. An analysis of 5’-noncoding sequences from 699 vertebrate messenger RNAs. Nucleic Acids Res 1987; 15(20): 8125–8148.

81. Gasteiger E, Hoogland C, Gattiker A, Duvaud S, Wilkins MR, Appel RD, et al. Protein identification and analysis tools in the ExPASy server. In: Walker JM (ed). The Proteomics Protocols Handbook. Humana Press, 2005, pp 571–607.

82. Robinson M, McCarthy D, Smyth G. edgeR: a Bioconductor package for differential expression analysis of digital gene expression data. Bioinformatics 2010; 26: 139–140.

